# Structure of full-length ERGIC-53 in complex with MCFD2 for cargo transport

**DOI:** 10.1101/2023.08.27.554937

**Authors:** Satoshi Watanabe, Yoshiaki Kise, Kento Yonezawa, Mariko Inoue, Nobutaka Shimizu, Osamu Nureki, Kenji Inaba

## Abstract

ERGIC-53 is a cargo receptor that promotes the transport of certain subsets of newly synthesized secretory proteins and membrane proteins from the endoplasmic reticulum (ER) to the Golgi apparatus (GA)^1,2^. Despite numerous structural and functional studies since its identification, the overall architecture and mechanism of action of this cargo receptor in its full-length form remain unclear. Here we present cryo-electron microscopy (cryo-EM) structures of full-length ERGIC-53 in complex with its functional partner MCFD2. These structures, in combination with SEC-MALS/SAXS analysis, reveal that ERGIC-53 exists as a homotetramer, not a homohexamer as previously suggested, and comprises a four-leaf clover-like head structure and a long stalk composed of three sets of four-helix coiled-coil followed by a transmembrane (TM) domain. The tetrameric head of ERGIC-53 consists of the vertically assembled carbohydrate recognition domains and the central four-helix coiled-coil. 3D variability analysis visualizes the globally flexible motion of the long stalk and local plasticity of the head region. Notably, MCFD2 has been found to possess a Zn^2+^ binding site in its N-terminal lid, which appears to modulate cargo binding. Altogether, unique mechanisms of regulated cargo capture and release by ERGIC-53 via the stalk bending and metal binding are proposed.

## Main

Secretory proteins and membrane proteins are newly synthesized in the endoplasmic reticulum (ER) and undergo oxidative folding and N-glycosylation with the help of luminal chaperones, glycosylation enzymes, and oxidoreductases^3,4^. A large portion of folded secretory proteins diffuse into the ER exit site, and are then transported into the Golgi apparatus (GA) by a bulk flow process^1^. However, subsets of secretory and membrane proteins are captured by cargo receptors in the ER, and subsequently transported to the GA in an accelerated manner^1^.

ERGIC-53 (also named LMAN1, P58) is a cargo receptor that captures certain specific secretory proteins^2,5^, including coagulation factor V (FV) and factor VIII (FVIII) ^6,7^, α1-antitrypsin^8–10^, cathepsin C^11^, cathepsin Z^12^, Mac-2 binding protein^13^, matrix metalloproteinase-9^14^ and IgM^15^. Mutations of the *ergic-53* gene cause a genetic bleeding disorder called combined deficiency of FV and FVIII (F5F8D)^7,16^. Another causative gene of F5F8D is the *multiple coagulation factor deficiency* 2 gene (MCFD2)^16,17^. MCFD2 is a small protein composed of an EF-hand domain with two Ca^2+^ binding sites, and functions as a binding partner of ERGIC-53 in a Ca^2+^-dependent manner^17–20^.

ERGIC-53 has also been reported to promote the ER-to-GA transport of the chaperone protein ERp44^21^ and several membrane proteins such as γ-aminobutyric acid type A receptors (GABA_A_Rs)^22^ and NDST1^23^. Furthermore, ERGIC-53 interacts with surface glycoproteins of infected RNA viruses and enhances their viral propagation^24,25^.

From the amino acid sequence, ERGIC-53 is predicted to be a single-pass transmembrane protein that consists of a luminal carbohydrate recognition domain (CRD), a long stalk domain, a transmembrane (TM) helix and a short cytoplasmic tail (CT)^5^. The CRD is structurally and functionally similar to a plant l-type lectin and binds to high-mannose type glycans of the target cargo proteins in a pH- and Ca^2+^-dependent manner^11,18,19,26,27^. The CT contains a C-terminal KKFF motif required for the ERGIC-53 cycling between the ER and GA^28–30^. The stalk domain is predicted to be composed of four long α-helices followed by a long loop and the single-pass TM helix. The three stalk α-helices except the most N-terminal helix are predicted to form a coiled-coil^31^. The long loop between the fourth stalk helix and TM domain contains two conserved Cys residues (Cys466 and Cys475), which form intermolecular disulfide bonds^31^.

After the discovery of ERGIC-53 in 1988, it was long believed that this cargo receptor existed as a covalent hexamer or covalent dimer via the interprotomer disulfide bonds between the stalk domain^32^. A subsequent study by Neve *et al*., however, demonstrated that ERGIC-53 existed in two forms, a covalent hexamer and a noncovalent hexamer, in which three of disulfide-bridged dimers of ERGIC-53 form the disulfide-linked hexamer (covalent hexamer) or are noncovalently assembled into the hexamer (noncovalent hexamer) ^33^. This preceding study also demonstrated that the cysteine mutant that lacks the intermolecular disulfide-bonds retained the hexameric state and was localized in the ERGIC as was wild-type ERGIC-53. It is thus suggested that the intermolecular disulfide bonds are not essential for the maintenance of its hexameric structure and cellular localization^33^.

Although the oligomeric states of ERGIC-53 were analyzed by biochemical approaches, more detailed structural and functional studies have been focused on the CRD and its complex with MCFD2^19,34–38^. Therefore, it still remained elusive how full-length ERGIC-53 forms the putative hexamer and works as a cargo receptor in this oligomeric state. Here, we present cryo-electron microscopy (cryo-EM) structures of full-length ERGIC-53 in complex with MCFD2. Contrary to the previous reports, the present structures, in combination with size-exclusion chromatography combined with multiangle light scattering (SEC-MALS) and with small angle X-ray scattering (SEC-SAXS) analysis, reveal a tetrameric rather than hexameric architecture, with a long flexible stalk domain. High resolution structures of its headpiece elucidate the molecular basis of the tetramer formation of ERGIC-53 and the zinc ion (Zn^2+^)-binding to the N-terminal region of MCFD2 for likely regulation of cargo binding and release. Altogether, the present study provides insights into the unique mechanism of cargo transport by ERGIC-53 using the long flexible stalk domain and the cargo-binding head region regulated by Zn^2+^.

### ERGIC-53 exists as a tetramer with a long extended structure

The previously proposed hexamer model of ERGIC-53 was based on PAGE analyses of crude purified samples^33^. To verify the oligomeric states of ERGIC-53 by more conclusive methods, we performed SEC-MALS analysis of purified full-length ERGIC-53. Recombinant full-length human ERGIC-53 was successfully purified to high purity from cumate-inducible stable HEK293T cell lines (Extended Data Fig. 1a). The monodisperse peak fractions from SEC elution contained two bands in nonreducing SDS-PAGE at ∼300 kDa and ∼110 kDa, which were presumed to be the covalent and noncovalent hexamers of ERGIC-53, respectively. However, SEC-MALS conjugated analysis showed that the molecular masses of purified ERGIC-53 and its complex with MCFD2 were determined to be 230 kDa and 280 kDa, respectively (Extended Data Fig. 1b, c and d). The determined molecular masses correspond to a homotetramer of ERGIC-53 alone (221.4 kDa) and a heterotetramer with four MCFD2 molecules (277 kDa), respectively. Thus, full-length ERGIC-53 likely exists as a mixture of covalent and noncovalent homotetramers, and its each protomer forms a 1:1 complex with MCFD2. The hexamer model (332 kDa) suggested by preceding works did not receive further experimental support.

To gain further insight into the overall structure of ERGIC-53 in solution, we next performed SEC-SAXS analysis (Extended Data Fig. 1e). The pair distance distribution function P(r) of full-length ERGIC-53 showed two separate peaks and estimated its maximum molecular dimension (*D*_max_) to be 347Å, suggesting that this cargo receptor adopts an overall dumbbell-like shape with an unusually long length (Extended Data Fig. 1f and 1h). Compared with isolated ERGIC-53, the ERGIC-53-MCFD2 complex displayed almost the same D_max_ and two slightly higher and sharper peaks in the P(r) function (Extended Data Fig. 1f and 1h), suggesting that in the complex, the corresponding two regions are more compact in conformation than those in isolated ERGIC-53. In the dimensionless Kratky plots of ERGIC-53 alone, the main peak was shifted to the larger qR_g_ side (∼ 3.5) and has a higher height (∼2.5) than those of typical globular proteins, suggesting that overall conformation of isolated ERGIC-53 is extended and in a highly asymmetric state ^39^. On the other hand, the complex showed not only a slightly narrower width of the peak but also lower values at the high qR_g_ region than ERGIC-53 alone, indicating that conformational flexibility or disorder was repressed upon complex formation with MCFD2 (Extended Data Fig. 1g). These results suggest that ERGIC-53 binds MCFD2 to adopt a more rigid or tightly folded structure than the isolate state.

### Cryo-EM structures of the head region of ERGIC-53 in complex with MCFD2

Based on the solution structure of the more rigid conformation of the ERGIC-53-MCFD2 complex, we performed cryo-EM analysis of full-length ERGIC-53 in complex with MCFD2. In the motion-corrected micrographs, long dumbbell-like particles of the complex were observed (Extended Data Fig 2, red square), consistent with the P(r) function obtained in the SEC-SAXS analysis. However, conventional autoparticle picking programs failed to recognize the full-length ERGIC-53 particles due to their unusual shapes. Instead, the autopicking programs recognized two globular portions in the dumbbell as separate particles (green circles in Extended Data Fig. 2). 2D classification of these two globular portions of the full-length particles showed clear 2D average images of the N-terminal tetrameric head of the complex and the C-terminal portion composed of the TM domain (including detergent micelle) and a part of the stalk domain, respectively (Extended Data Fig. 3a). Consequently, we determined the cryo-EM structures of the head region of full-length ERGIC-53 complexed with MCFD2 in two forms (forms A and B) at 2.53 Å and 2.59 Å resolutions, respectively (Extended Data Fig. 3a, b). On the other hand, only low-resolution EM maps were obtained for the C-terminal portion, probably due to its high flexibility and heterogeneity in the shape of detergent micelles.

The present cryo-EM analysis reveals that the overall structure of the assembled head is composed of four ERGIC-53 protomers and four MCFD2 molecules, confirming that ERGIC-53 exists as the tetramer (Fig. 1a, b). Each head region consists of the CRD (residues 31-268), stalk helices 1 and 2 (S-H1 and S-H2), and loops connecting these domains (Fig. 1c, left). The tetrameric structure is stabilized by a central four-helix coiled coil composed of the long S-H2 from each protomer (residues 326-367) (Fig. 1b). The local resolution of S-H1 and the loops between CRD and S-H1, and between S-H1 and S-H2 are relatively low, suggesting the high flexibility of these regions (Extended Data Fig. 3c). MCFD2 binds the CRD at a 1:1 molar ratio as observed in the crystal structures of the CRD-MCFD2 complex^34–37^. In addition, the S-H1 weakly associates with a part of MCFD2 (Fig. 1c and Extended Data Fig. 4a, inset 3), likely affecting the conformation of its N-terminal region (also see below).

**Fig. 1.**
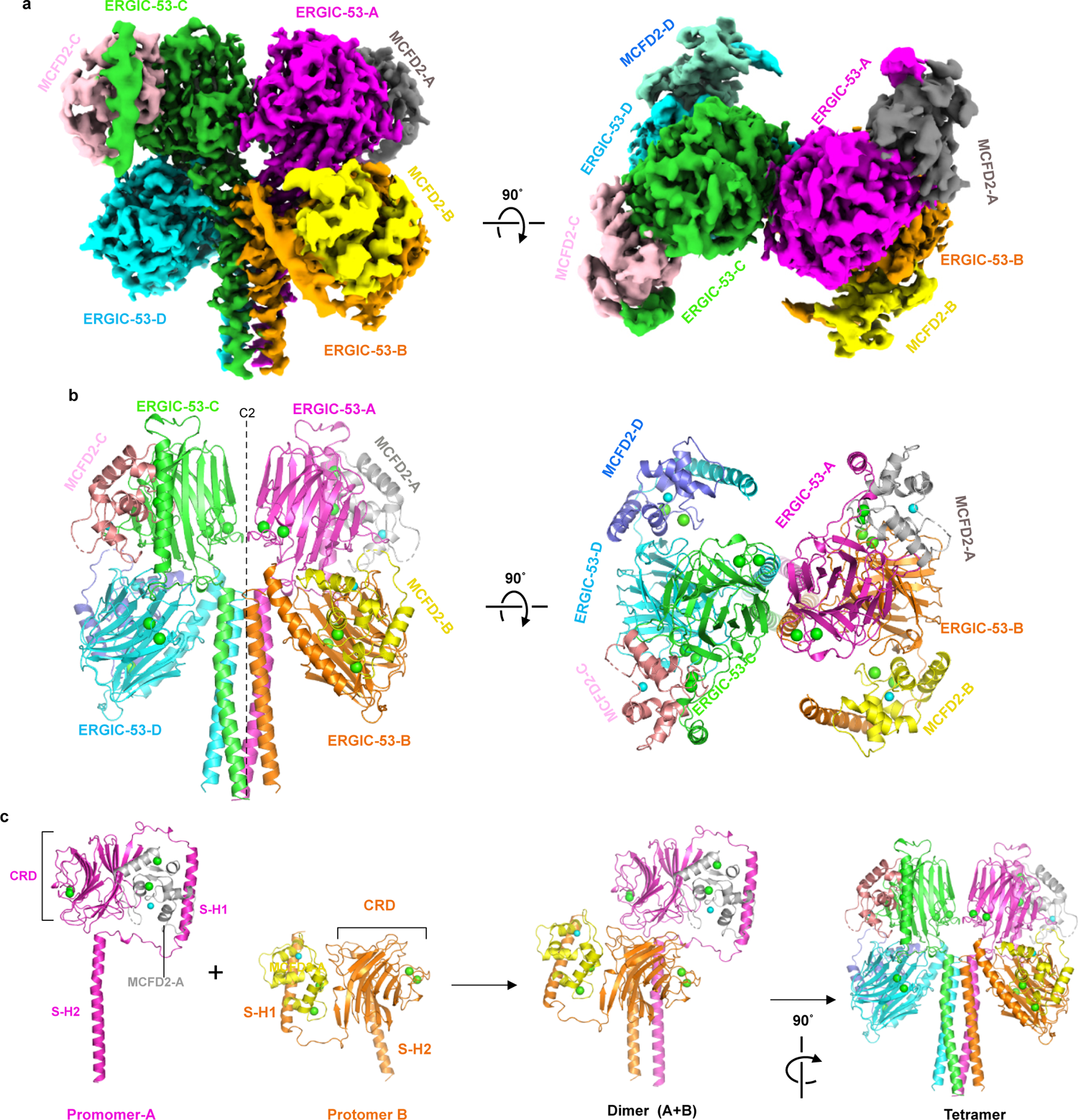
Cryo-EM structures of the head region of ERGIC-53 in complex with MCFD2 a) Side and top views of the cryo-EM map of the head region of ERGIC-53 with MCFD2. Four ERGIC53 protomers and four MCFD molecules are shown in magenta (ERGIC-53 protomer A), orange (protomer B), green (protomer C), cyan (protomer D), gray (MCFD2-A), yellow (MCFD2-B), pink (MCFD2-C), and light blue (MCFD2-D), respectively. b) Ribbon diagram of the structure of the tetrameric head region of the ERGIC-53. The four ERGIC-53 protomers and four MCFD2 molecules are shown in the same color as in 1a. Green and cyan spheres represent calcium and zinc ions, respectively. c) Details of the tetrameric head formation. The left two panels show structures of the head region of the ERGIC-53 protomers in the upper CRD configuration (protomer A) and lower-CRD configuration (protomer B).

### CRD head assembly

Although ERGIC-53 forms the homotetramer, the four CRDs are not related by a four-fold axis, but arranged in a C2 symmetry. Two CRDs are located in the upper layer, whereas the other two lie in the lower layer (Fig. 1b). In the protomers with the upper CRD (protomers A and C), the head region adopts a “Hisyaku (Japanese ladle)”-like shape, in which both CRD and S-H1 are positioned above the S-H2 (Fig. 1c). MCFD2 is accommodated in the pocket formed by CRD, S-H1 and two long loops between CRD and S-H1 and between S-H1 and S-H2. In the protomers with the lower CRD (protomers B and D), CRD and S-H1 are rotated by approximately 180° around the SH1-SH2 loop, and are located below the top of S-H2. The upper-CRD protomer vertically associates with the lower CRD protomer to form a dimer, and then the two dimers are further assembled to form the four-leaf clover-like tetramer (Fig. 1b, c).

In the assembled head region, two different CRD-CRD interfaces are formed: a vertical interface (interface-V) between the vertically assembled CRDs (between protomers A and B and between protomers C and D) (Extended Data Fig. 4a) and the other interface (interface-H) between the horizontally assembled upper CRDs (between protomers A and C) (Extended Data Fig. 4b). Interface-V is formed by van der Waals contacts between residues in the β-strands and their flanking loops of each CRD (Extended Data Fig. 4a, inset 1). A similar interface is also observed in the crystal packing of the CRD-MCFD2 complex (PDB code: 3WHU, 4YGE, etc.). The lower CRD (protomer B) also interacts with S-H2 from the counter protomer (protomer A) (Extended Data Fig. 4a, inset 2). On the other hand, interface-H is formed by only four hydrogen bonds including that between Arg192 and Gln191’ and two van der Waals contacts between the CRDs (Extended Data Fig. 4b).

As described above, two different forms (A and B) of the head region are determined, in which the positions and orientations of the CRDs relative to the S-H2 are different. The superimposition on the central S-H2 coiled coil demonstrates that during the conversion from form A to form B, the upper and lower CRDs rotate by 4.8° as a rigid body (Extended Data Fig. 4c). Consequently, interface-H in form B is formed by interactions between different residues, such as van der Waals contact between Gln191 and Glu228’ (Extended Data Fig. 4b right), while interface-V is almost maintained. These results suggest that interface -V involves tight binding, whereas interface-H is variable. As a result, the vertically assembled pair of CRDs can rotate to a certain extent (by 4.8°) as a rigid body.

### Overall architecture of full-length ERGIC-53 with MCFD2

Next, we endeavored to determine the whole structure of the full-length ERGIC-53-MCFD2 complex. Manual picking of full-length particles was first conducted based on the positions of the head and TM regions. An initial picking model was trained by Topaz^40^ using particles selected from 2D classification, and the picking model was further improved by repeating Topaz picking, 2D classification and Topaz training. Finally, Topaz successfully recognized the full-length particles and enabled us to perform single particle analysis of full-length ERGIC-53 (Extended Data Fig. 5a).

2D classification of the collected full-length particles generated clear 2D average images of their four-leaf clover-like structures consisting of the head, long stalk and TM regions (Fig. 2a). Some particles adopt nearly straight conformations, while others possess stalks largely bent at various angles. These results suggest that the stalk region of ERGIC-53 is highly flexible.

**Fig. 2.**
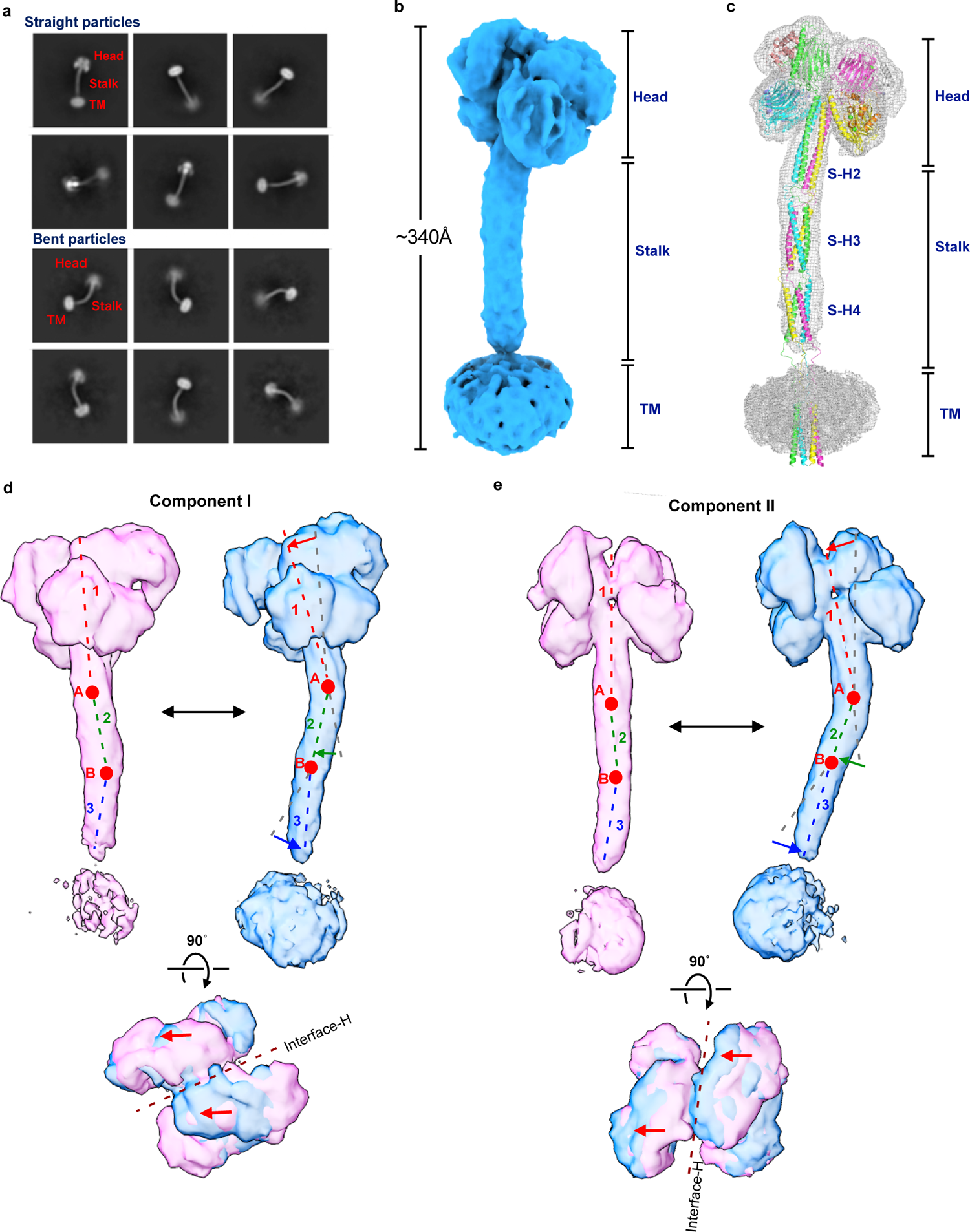
Cryo-EM structure of full-length ERGIC-53 in complex with MCFD2 a) Representative 2D class average images of the full-length particles. Upper and lower panels represent straight and bent particles, respectively. b) Cryo-EM map of full-length ERGIC-53 in complex with MCFD2. c) The overall architecture of full-length ERGIC-53 overlaid in the EM map shown in (b). d) Results of 3DVA of the full-length ERGIC-53 with two variability components I and II. The 3D EM maps of the first (pink) and last frames (blue) of continuous conformational changes generated by 3DVA are displayed. Broken lines represent the center of three segments. Red circles represent hinges between the segments 1, 2, and 3. The lower panels show top views of the two maps superimposed on each other. The brown broken line represents the interface-H between the upper CRDs.

By using straight particles, the cryo-EM structure of full-length ERGIC-53 in complex with MCFD2 was determined at 6.8 Å resolution (Fig. 2b). The maximum length of the full-length ERGIC-53 structure is approximately 340Å, which is almost comparable to the *D*_max_ value (347 Å) obtained by SAXS analysis (Extended Data Fig. 1f). Based on the obtained cryo-EM map, we built a structure model of full-length ERGIC-53 by using AlphaFold multimer^41^ (Fig. 2c). Similar to S-H2, the stalk helices 3 (S-H3) and 4 (S-H4) also appear to form a four-helix coiled-coil. Thus, the three sets of long four-helix coiled-coils are vertically connected by relatively long loops to form the stable but flexible tetramer. Although the TM helices were not well resolved in the current cryo-EM map, they are also predicted to form a four-helix bundle and be vertically connected to S-H4. As a result, full-length ERGIC-53 adopts a slender overall architecture. To our knowledge, the molecular height of full-length ERGIC-53 is taller than that of any other membrane protein of known structure, including the largest Ca^2+^ channel ryanodine receptor RyRs, V-type ATPase, and spike protein from COVID-19 (Extended Data Fig.5c). To assess the structural contribution of MCFD2 to the overall architecture of ERGIC-53, we also attempted the cryo-EM analysis of full-length ERGIC-53 alone. While 2D average images showed features of the C-terminal portion including SH-4 and the TM domain, clear 2D images were not observed for the tetrameric head regions (Extended Data Fig.6a). In the full-length particle analysis by using Topaz, the head regions were highly blurred after 2D averaging (Extended Data Fig.6b). Consistently, our SEC-SAXS analysis suggests that the structure of full-length ERGIC-53 is more flexible in the absence of MCFD2 (Extended Data Fig. 1f, g). Thus, MCFD2 binding seems likely to stabilize the structure of the tetrameric head of ERGIC-53.

### Detailed molecular motions of full-length ERGIC-53

To gain insight into the molecular motions of full-length ERGIC-53, 3D variability analysis (3DVA) was performed in cryoSPRAC^42^, resulting in identification of two different continuous motions of this cargo receptor (components I and II) (Fig. 2d,e, Supplementary movie S1). 3DVA suggests that the entire structure of ERGIC-53 can be divided into three rigid-body segments, which correspond to the head region (segment 1), S-H3 (segment 2), and S-H4 (segment 3), respectively, followed by the C-terminal TM helix. The hinges between the three segments correspond to loops between S-H2 and S-H3 (hinge A) and between S-H3 and S-H4 (hinge B). In component I, the segments 1 and 2 undergo a slight bending and stretching motion around hinge A. Segment 3 also undergoes a swing movement, rotating about hinge B in the opposite direction to segments 1 and 2 (Fig. 2d). The rotation directions of these three segments are nearly parallel to the interface-H between the upper CRDs (Fig. 2d bottom). The density of the TM domain was largely variable during the conversion from the first to last frames in this component, suggesting that the TM domain also swings around the loop flanked by S-H4 and the TM domain (i.e., the third hinge) (Fig. 2d). In the component II, the segments 1-3 undergo similar bending and stretching movements through the two hinges, in which their rotation directions are perpendicular to interface-H (Fig 2e, bottom). In addition, other conformations with even more greatly bent stalks were also observed in the 2D class-average images of the intact particles (Fig. 2a lower panels). These results suggest that full-length ERGIC-53 continuously adopts diverse conformations by bending stalk and TM helices at the three hinges.

### Molecular motion of the head region in full-length ERGIC-53

As mentioned above, the two different structures of the head region suggest its conformational variability (Extended Data Fig. 4c). To further assess the conformational variability of the head region, 3DVA was also performed by focusing exclusively on this region (Extended Fig. 7a, Supplementary movie S2). As observed in forms A and B (Extended Data Fig. 4c), component I^head^ shows a swing movement of the CRDs, in which the CRD dimer between upper and lower protomers shows a side-to-side rotation as a rigid body dimer (Extended Data Fig. 7a left, movie S2 left). The rotation direction of the dimer is nearly parallel to interface-H (side-to-side rotation) (Extended Fig. 7a lower left).

In component II^head^, two CRD dimers rotate along a vertical axis (i.e. nearly parallel to the central coiled-coil) in opposite directions (referred to as twisting motion) (Extended Fig. 7a lower right, movie S2 right). Based on a clustering analysis of the 3DVA of the head region, four different substate structures (A-D) were reconstructed at 3.29∼3.51 Å resolution (Extended Data Fig. 6b, c). Superimposition of the central coiled coil helices from the substates A and B reveals that the upper and lower CRD dimers are rotated by 6.2° as a rigid body, which is a larger rotation between form A and form B described above, suggesting that form A and form B represent more averaged structures of the head region individually (Extended Data Fig. 7d, vs. 4c). In the superposition of substate structures C and D, SH1 and MCFD2 in the upper layers and those in the lower layers rotate by 2.5° and 1.9°, respectively, breaking the C2 symmetry of the head region (Extended Data Fig. 7e). These rotations seem to be ascribed to the flexible loops between the CRD and S-H1 and between S-H1 and S-H2. Thus, the two loops within the head region likely generate the plasticity of the head region, allowing the CRD-MCFD2 complex to fluctuate continuously in the tetrameric structure.

### Local plasticity of the head region independent of the global stalk bending

To investigate the linkage between the local plasticity of the head region and the global stalk bending, we constructed an ERGIC-53 1′H34 mutant in which S-H3 and S-H4 within the stalk were deleted, and determined its cryo-EM structure at 3.5Å resolution (Extended Data Fig. 8a). The ΔH34 mutant retains a tetrameric structure with the assembled head region, similar to wild-type. Additionally, the TM helices of the mutant were not well resolved in the density map, as was the case for wild-type. 3DVA of this deletion mutant revealed its three different motions (Extended Data Fig. 8b), Supplementary movie S3). In the component I^ΔΗ34^, the TM domain shows a large swing movement, while the head region remains static (Extended Data Fig. 8b, left, movie S3 left). On the other hand, components II^ΔΗ34^ and III^ΔH34^ show the side-to-side rotation and twisting motion of the CRD dimers, respectively (Extended Data Fig. 8b center and right, and Supplementary movie S3 center and right), as observed in the 3DVA of the head region of full-length ERGIC-53. The TM domain shows only marginal rotation in either component II^ΔH34^or III^ΔΗ34^. Thus, the CRD dimers in the head region appear to move separately from the region below the S-H2, suggesting that there is no linkage between the head-domain plasticity and the stalk-domain bending.

### Visualization of an N-terminal segment with a zinc binding site in MCFD2

In the previously reported crystal structures of MCFD2 alone and its complex with CRD ^20,34–37^, the N-terminal region of MCFD2 (residues 27-66) and a loop between α3 and α4 helices (residues 100-111) were disordered and hence invisible (Extended Data Fig. 9a). However, the present cryo-EM map displays significant density for the regions preceding the α2 helix (Fig. 3a). In this context, AlphaFold2 prediction^41^ suggests that the disordered N-terminal region contains two short α-helices (α1-A, and α1-B) (Extended Data Fig. 9b). Although we had difficulty in *de novo* modelling of the MCFD2 N-terminal region from the present map, the predicted N-terminal α1-A, and α1-B short helices and a loop between the α1-B and α2 helices are nicely fitted onto the observed density map (Fig. 3a). The difference in the maps of MCFD2 obtained by the crystal and cryo-EM analyses is likely due to the presence of S-H1 of ERGIC-53 in the cryo-EM sample. Indeed, this helical segment interacts with the N-terminal region of MCFD2, seemingly restricting its conformational flexibility (Fig. 2c, Extended Data Fig. 4a).

**Fig. 3.**
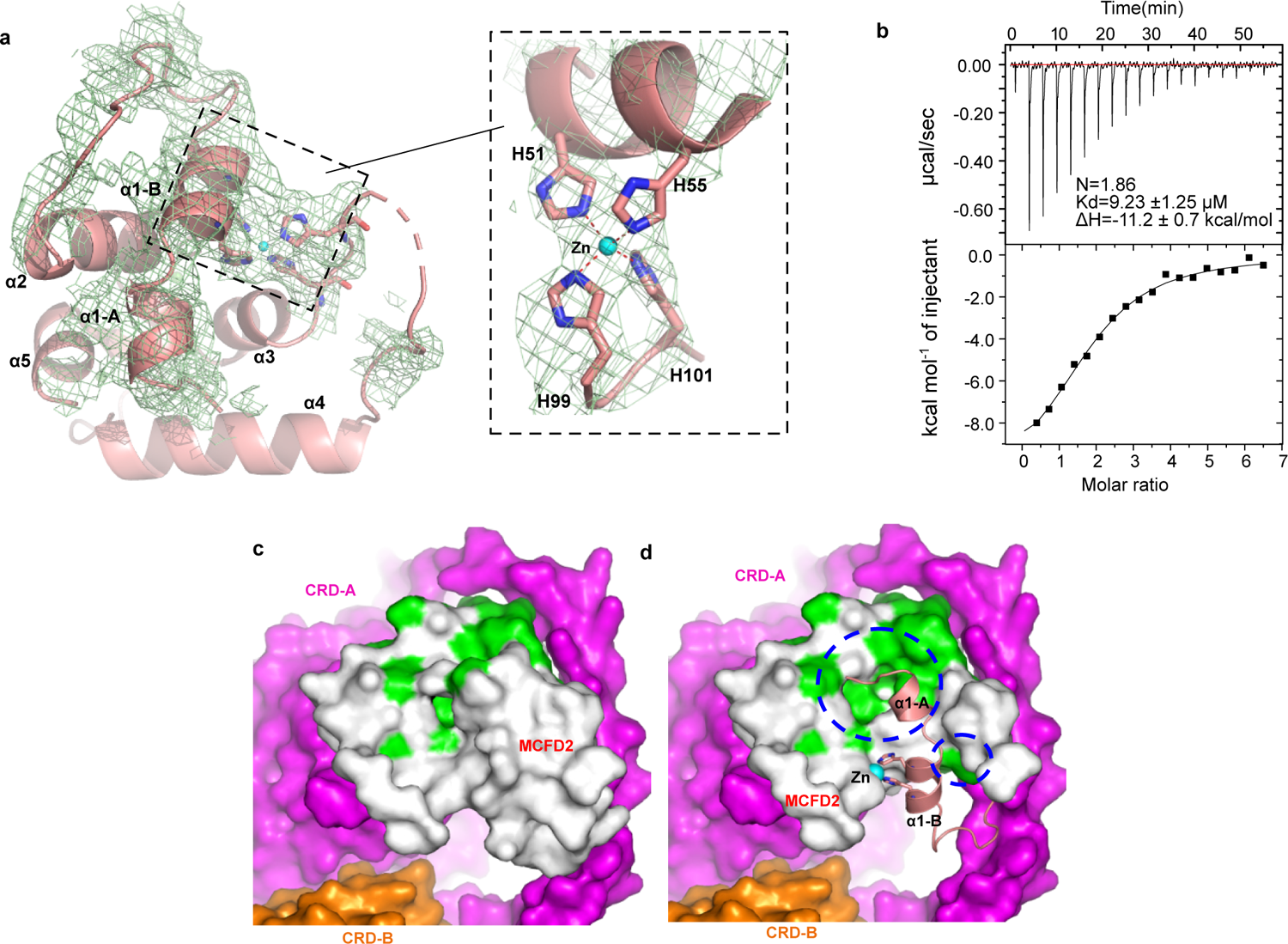
Visualized N-terminal segment of MCFD2 contains a Zn2+ binding site a) The overall structure of MCFD2 bound to full-length ERGIC-53. The EM density map around the region that is missing in the previous crystal structures is shown in a light green mesh. The inset shows a close-up view of the His cluster, showing that Zn^2+^ is probably coordinated by four His residues. b) ITC raw data (upper) and binding isotherm data (lower) for titration of ZnCl_2_ (500 µM) into the EDTA-treated MCFD2 sample. Errors represent fitting residuals (standard deviation). c, d) Comparison of FV/FVIII binding sites in MCFD2 with (c) or without (d) the N-terminal lid. MCFD2 (white) and two CRDs (magenta and orange) are shown in the surface representation. Previously identified FV/FVIII binding sites in MCFD2 are colored in green. Blue dashed circles indicate the regions of the FV/FVIII binding site masked by the N-terminal lid.

In the updated MCFD2 structure, four conserved His residues (His51, His55, His99 and His101) are clustered at the α1-B helix and a loοp between the *α*3 and α4 helices (Fig. 3a, Extended Data Fig. 9d). Notably, the extra density is clearly seen at the center of the His cluster in the present cryo-EM map (Fig. 3a inset), suggesting that a metal ion, most likely a zinc ion, is coordinated by four histidine residues. In support of this, a colorimetric assay with 4-(2-pyridylazo) resorcinol (PAR), a chromophoric chelator for divalent metal ions, revealed that purified MCFD2 contained significant amounts of divalent metal ions such as Zn^2+^ (Extended Data Fig 9c). Consistently, our isothermal titration calorimetry (ITC) analysis revealed that an EDTA-treated MCFD2 sample can bind up to two molar equivalents of Zn^2+^ with micromolar affinity (Fig. 3b). Thus, MCFD2 has been shown to be a zinc-binding protein with the conserved His cluster.

A previous NMR study revealed that the EF hand helices of MCFD2 are involved in the interaction with FV/FVIII^43^. In the present cryo-EM structures, the reported FV/FVIII binding site in MCFD2 is not fully exposed to the solvent but rather shielded by its N-terminal short helices (Fig. 3c, d). The N-terminal segment of MCFD2 appears to be hooked by the Zn^2+^-bound His cluster formed between the α1-B helix and α3-α4 loop (Fig 3a). These structural features raise a possibility that Zn^2+^ binding to the His cluster may prevent the cargo binding to the head of ERGIC-53.

To explore how Zn^2+^ affects the cargo transport, we performed an FV secretion assay using WT and ERGIC-53 KO 293T cells with or without Zn^2+^ supplementation. Intracellular Zn^2+^ concentration is known to be affected by extracellular Zn^2+^ levels in culture medium^44^. When 10 µM ZnCl_2_ was added in the serum-free Expi293 medium, FV secretion from WT cells was slightly but significantly prevented (Extended Data Dig. 9e, left). In contrast, the ERGIC-53 KO cells hardly displayed the Zn^2+^-induced prevention of FV secretion (Extended Data fig 9e, right). Altogether, these results suggest that the cargo transport mediated by ERGIC-53/MCFD2 can be inhibited by Zn^2+^, consistent with the present observation that the cargo-binding site of ERGIC-53 is masked by the N-terminal segment of “Zn^2+^-bound” MCFD2.

### Physiological significance of the long stalk and head assembly

Our cryo-EM structures reveal that full-length ERGIC-53 adopts the four-leaf clover-like structure with the unusually long and flexible stalk. To examine the physiological roles of the long stalk in the cargo transport, rescue experiments were performed by monitoring FV and alpha-anti-trypsin (AAT) secretion under coexpression of the cargo proteins and ERGIC-53 WT or its stalk-deletion mutant in the ERGIC-53 KO cells (Fig. 4a). Upon cotransfection, the expression levels of WT and its mutant were comparable to that of endogenous ERGIC-53 in HEK293T cells (Extended data Figs. 11 a, c). As expected, the coexpression of ERGIC-53 WT enhanced the secretion of FV and AAT, compared to the empty vector (Fig 4b, c, Extended Data Fig. 11b, e). Similarly, expression of the 1′H34 mutant with a much shorter stalk (Fig 4a) rescued the FV section at a level comparable to WT (Fig. 4b). A similar result was obtained for the AAT secretion (Fig. 4c). Thus, deletions of the S-H3 and S-H4 only marginally affected the cargo transport ability of ERGIC-53. To verify the validity of our ERGIC53-mediated cargo secretion assay, we also investigated the FV and AAT secretion from the ERGIC-53 KO cells expressing ERGIC-53 W67S, a missense mutant deficient in both MCFD2 and mannose bindings^45^. As reported previously^45,46^, this negative control mutant failed to rescue the FV secretion while the AAT secretion was partially rescued by this mutant (Fig. 4b, c) probably because AAT is an MCFD2-independent cargo^47^. Collectively, it can be interpreted that the shorter version of ERGIC-53 (ΔH34) still retains the ability to enhance the secretion of FV and AAT although the distance between the head and TM domain and the flexibility of the stalk are substantially reduced compared to those of wild-type.

**Fig. 4.**
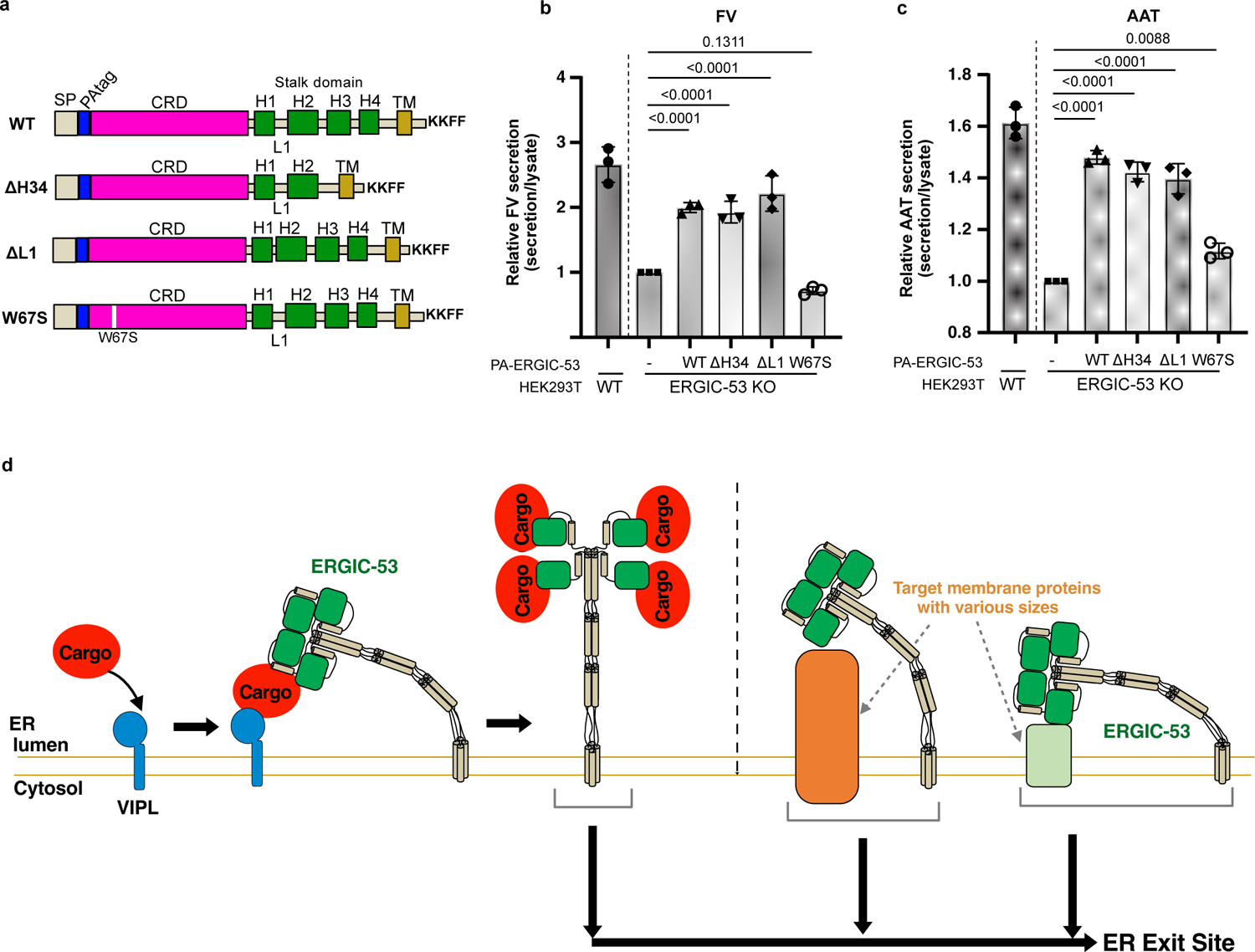
Role of the long stalk and the head assembly in cargo transport a) Schematic representation of the ERGIC-53 mutants used for the rescue experiments. b, c) Rescue experiments of FV-HiBiT (b) and AAT-HiBiT (c) secretion by coexpression of ERGIC-53 or its mutants in ERGIC-53 KO cells. The amounts of secreted FV/AAT relative to those of intracellular FV/AAT were quantified with a commercially available HiBiT-tagged protein detection reagent and normalized to those from the KO cells transfected with empty vector. Dots represent the individual data points. Error represents the standard error of the mean (N=3 biological replicates). Statistical significance was measured with one-way ANOVA followed by Dunnett’s test. d) A working model of cargo capture by ERGIC-53.

Additionally, to examine the physiological significance of the head region assembly for cargo transport, we constructed a mutant lacking the L1 loop between S-H1 and S-H2 (residue Gln313 to Glu 322) (ΔL1 mutant) (Fig. 4a, Extended Data Fig. 10c). In the head region, two S-H1s of protomers A and B are located ∼65 Å apart by the L1 loops to form the vertically assembled CRD dimers. (Extended Data Fig. 10b). Hence, deletion of L1 loop is predicted to disrupt the interface-V between the upper and lower CRDs, as shown in the AF2 predicted model (Extended Data Fig. 10c and d). The 1′L1 mutant displayed a similar migration pattern to the WT on a nonreducing SDS gel (Extended Data Fig. 11a lane 5), suggesting that this mutant also exists as the mixture of covalent and noncovalent tetramers, likely formed through the central four-helix coiled coils. Coexpression of the 1′L1 mutant also rescued the FV and AAT secretion at a level similar to that of the WT. These results suggest that the head region assembly is not an essential factor for the efficient transport of soluble cargo proteins by ERGIC-53 and MCFD2.

## Discussion

The present cryo-EM analysis reveals the tetrameric slender architecture of full-length ERGIC-53 in complex with MCFD2. The long stalk of ERGIC-53 is composed of three sets of four-helix coiled coil that are vertically aligned. Full-length ERGIC-53 can adopt various conformations including straight and largely bent conformations by bending the stalk domain. The CRDs within the head region also fluctuate independently of the stalk bending. These flexible motions of ERGIC-53 are most likely generated by the loops that function as hinges between the CRD, stalk helices and TM domain.

ERGIC-53 has been proposed to cooperate with other luminal L-type lectins such as VIP36(LMAN2), VIPL(LMAN2L), and ERGL (LMAN1L), to maintain the homeostasis of secretory cargo proteins^26,48,49^. In the proposed model, properly folded cargo proteins with M9 glycans are first recognized by VIPL, and then transferred to ERGIC-53. VIPL consists of a CRD and a single TM but lacks a stalk region. ERGIC-53 presumably receives cargo proteins from the small VIPL by utilizing the largely bent long stalk (Fig. 4d left). In this context, it is reasonable that the ΔH34 mutant with the much shorter stalk still retains the cargo transport activity at a similar level to WT. Our rescue experiments also revealed that the head assembly is not essential for the efficient cargo transport. Given the docking model of dimannose bound to the CRD based on the previous crystal structure (4YGE), the 1-OH of the Man (4) moiety of Man_9_(GlcNAc)_2_ glycan is located near the L1 loop between S-H1 and S-H2 (Extended Data Fig. 9f). In the tetrameric head, the L1 loop may interfere with the accommodation of N-glycans of cargo proteins, raising the possibility that the formation of the head assembly rather promotes the release of cargo proteins. Conversely, upon cargo binding, the assembled CRDs may disassemble, allowing each CRD to capture a target cargo protein in concert with MCFD2.

In addition to soluble secretory proteins, ERGIC-53 is known to facilitate the transport of certain specific membrane proteins including neuroreceptors such as GABA_A_R in neuronal cells^22^ and several Golgi resident membrane proteins such as GOLGI5M, NDST1 and FKRP^23^. ERGIC-53 also associates with surface glycoproteins of arenavirus, hantavirus, coronavirus and hepatitis B virus^24,25^. Given that these target membrane proteins vary in their size and structures, the conformational flexibility of the stalk domain and the local rotation of the CRDs appear to help adjust the height and position of the head region of this cargo receptor to those membrane proteins (Fig. 4d right). Notably, although the CRD of ERGIC-53 is an essential element for the transport of these membrane proteins, its glycan binding ability is dispensable for the interaction with them ^24,25^. Collectively, ERGIC-53 is likely to recognize and bind target membrane proteins differently from soluble proteins.

Here, MCFD2 has been found to be a zinc binding protein. In the Zn^2+^-bound structure, the cargo-binding site in MCFD2 is masked by its N-terminal helix, and this local structure is probably stabilized by the Zn^2+^-bound His cluster (Fig. 3a). The relatively low local resolution of the N-terminal region of MCFD2 may suggest that this segment is readily moved away upon cargo binding. Thus, the N-terminal helix of MCFD2 appears to function as a lid to regulate the cargo binding/release in a Zn^2+^-dependent manner (Extended Data Fig. 12). It has been presumed that ERGIC-53 releases cargo proteins in the GA with lower Ca^2+^ concentration and lower pH^11,18,19,26,27^. In contrast to Ca^2+^, the labile Zn^2+^ concentration has been reported to be much higher in the GA than in the ER and other organelles^50–52^. Accordingly, these findings suggest that Zn^2+^ binds transiently to MCFD2 in the GA and thereby promotes the cargo release by closing its N-terminal lid. The proposed working hypothesis of Zn^2+^-dependent regulation of the ERGIC-53 MCFD2 system is relevant to our recent finding that Zn^2+^ is vital to protein quality control mediated by ERp44 in the early secretory pathway (ESP) ^51,53^. In sharp contrast to the ERGIC-53-MCFD2 system, Zn^2+^ enhances the complex formation between ERp44 and its client proteins in the GA, whereas ERp44 releases its clients in the ER at much lower Zn^2+^ concentrations^51,53^. Future studies will further elucidate the physiological roles of Ca^2+^, Zn^2+^ and other metal ions in protein homeostasis mediated by various molecular chaperones and cargo receptors in the ESP.

As an additional note, the revealed slender structure of full-length ERGIC-53 seems highly relevant to C-type lectins, which are also composed of a CRD, long stalk, and TM domain. Many of the C-type lectin family proteins are localized on the cell surface and function as receptors for a broad range of ligands, including pathogens^54,55^. Intriguingly, a recent study reported that in dendritic cells and relevant cells, a fraction of ERGIC-53 is located on the cell surface and functions as a receptor for house dust mite (HDR) allergens^56^. The molecular height of ERGIC-53, which is comparable to those of the C-type lectins, is likely advantageous for this receptor to capture target HDR and other potential ligands, without substantial interference from surrounding C-type lectins on the cell surfaces. It is still difficult to visualize cell surface receptors such as full-length C-type lectins due to their unusually long stalks and large flexibility. Nonetheless, as shown here, cryo-EM single particle analysis can visualize the overall structures of the flexible receptor not only in the 2D class average images but also in 3D reconstruction at a middle resolution that enables the reliable identification of each domain. Flexibility analysis of the EM data such as 3DVA can visualize their global motion. Further structural analysis of long flexible receptors will be feasible and provide a deep understanding of their structures and mechanisms of action.

## Methods

### Cell cultures and Plasmids

The cDNA of human ERGIC-53 was subcloned from a cDNA library from HeLa cells and inserted into the pcDNA3.1 vector (Invitrogen). Subsequently, the ERIGIC-53 gene with a C-terminal PA tag was inserted into a PiggyBac Cumate Switch Inducible vector (System Bioscience). The pcDNA-SP-PA-ERGIC-53 plasmid was generated using the In-Fusion cloning kit (Taraka) with PCR fragments from the pcDNA3.1-ERGIC-53-PA plasmid. Mutants of ERGIC-53 were constructed by PCR-based site-directed mutagenesis. A PCR fragment of the ΔΗ34-mutant with a C-terminal FLAG was also inserted into a pEMmulti-puro vector (Fijifilm-Wako).

The cDNA of human MCFD2 (IRAK026G06) was provided by the RIKEN BRC through the National BioResource Project of the MEXT/AMED, Japan. The lumen domain of MCFD2 (residues 27-146) was subcloned into the pET15b vector with an N-terminal histidine tag. The cDNA of human Factor V (FV) (FXC01788) was purchased from Kazusa DNA Research Institute. The secreted region of FV (residues 29-2224) was inserted into the phlsec-vector (Addgene, 72348) with the C-terminal HiBiT-Tag (Promega), using the In-Fusion cloning kit. A codon-optimized cDNA fragment of the secreted region of human AAT was synthesized (eurofin) and inserted into the phlsec-vector with the C-terminal HiBiT-Tag.

Human embryonic kidney (HEK) 293T cells were purchased from ATCC (American Type Culture Collection. Human ERGIC-53 (LMAN1) KO 293T cell lines were purchased from Abcam (ab266248).

### Protein expression and purification

The PiggyBac-cumate-ERGIC-53-PA tag plasmid along with the Super PiggyBac Transposase Expression Vector (System Bioscience) was transfected to generate a stable HEK293T cell line with cumate inducible ERGIC-53 expression. The established stable cells were cultured in Dulbecco’s modified Eagle’s medium (DMEM, Nacalai Tesque) supplemented with 4∼5% fetal calf serum (FCS; Thermo Fisher Scientific, or Nichirei Biosciences Inc) and 1% penicillin-streptomycin mixed solution (Nacalai Tesque) under 5% CO_2_ at 37°C. When cells were grown to 70∼80% confluency in 15 cm dishes, the medium was replaced with serum-free DMEM supplemented with 6 mM sodium butyrate (Nacalai Tesque). After 6 h incubation at 37°C, expression of ERGIC-53 was induced by the addition of cumate (240 µg/ml) and phorbol 12-myristate 13-acetate (PMA, 50 nM), and then cells were cultured at 30 °C in 5% CO_2_ for another 2 days before harvesting. The cells were washed with PBS and stored at −80°C. The pEB-multi-puro-ERGIC-53 H34-FLAG was transfected to generate a stable HEK293T cell line for expression of the ERGIC-53 ΔH34-FLAG mutant. The established stable cells were cultured in DMEM supplemented with 4∼5% FCS and 1% penicillin-streptomycin mixed solution (Nacalai Tesque) under 5% CO_2_ at 37°C. When cells were grown to 70∼80 % confluency in 15 cm dishes, sodium butyrate (6 mM) and PMA (50 nM) were added and then cells were cultured at 30°C in 5% CO_2_ for another 2 days before harvesting. The cells were washed with PBS and stored at −80°C.

All purification procedures were conducted at 4°C or on ice. Cell pellets expressing ERGIC-53-PA from 3 liter of medium or those expressing the ERGIC-53 ΔH34 mutant from 1 litter of medium were resuspended in buffer A (20 mM Tris-HCl pH 7.5, 150 mM NaCl, 5 mM CaCl_2_, 10% glycerol) supplemented with 5 mM MgCl_2_ and 1x protein inhibitor cocktail (Nacalai Tesque) and ∼50 µg/ml DNase I (Fujifilm-Wako). The cells were broken by brief sonication and were solubilized by the addition of 1% (v/v) n-dodecyl-β-D-maltoside (DDM; Nacalai Tesque) for 2 h. The solubilized fraction was clarified by ultracentrifugation (Hitachi CS100FNX, 150,000g, 20 min) and then mixed with 7.5 ml anti-PA-tag antibody beads (Fujifilm-Wako) for 2 h (for ERGIC-53-PA) or 1 ml Anti-DYDDDK tag antibody beads (MBL) for 2 h (for the ERGIC-53 ΔΗ34 mutant) The beads were transferred into a gravity flow column (Econo-column, Bio-Rad) and washed with 20 CV of buffer A supplemented with 0.02% (w/v) GDN (Anatrace). The bound proteins of ERGIC-53 WT were eluted with overnight incubation of 2 CV elution buffer (buffer A supplemented with 0.02% GDN, and 0.2 mg/ml PA peptide), followed by repeating 5 min incubation with 1CV elution buffer. For the ΔΗ34 mutant, the bound proteins were eluted by repeating the 5 min incubation of 1 CV elution buffer (buffer A supplemented with 0.02% GDN and 0.1 mg/ml DYKDDDDK peptide). The eluted protein was concentrated using Amicon Ultra-15 centrifugal filters (100 kDa cut-off, Merck-Millipore). After treatment with 1 mM diamide for 15 min to completely oxidize the protein, the concentrated protein was applied onto a Superose 6 Increase 10/300 GL column (GE healthcare) equilibrated with SEC buffer (20 mM Tris-HCl pH 7.5, 150 mM NaCl, 10 mM CaCl_2_ and 0.02% GDN). For preparation of the ERGIC-53-MCFD2 complex, the ERGIC-53 sample was mixed with excess purified MCFD2 (described below) for 10 min, followed by diamide treatment for 15 min and then the complex sample was injected into the same column. The peak fractions were concentrated using Amicon-Ultra-4 and −0.5 centrifugal filters (100 kDa cut-off) for cryo-EM sample.

The pet15b plasmid of MCFD2 was introduced into E-cos Escherichia coli BL21 (DE3) (Nippon gene). Gene expression was induced with 0.1 mM isopropyl β-D-1-thiogalactopyranoside (Fujifilm Wako) at an OD600 of ∼0.5, with further cultivation for 24 h at 293 K before harvesting. Cells were washed with 50 mM Tris-HCl pH 8.5 and then stored at −80°C. The cell pellet was resuspended in buffer A (50 mM Tris-HCl pH 8.0, 300 mM NaCl, 5 mM CaCl_2_) supplemented with protease inhibitor cocktail (Nacalai Tesque) and was sonicated on ice for 5 min. After removal of cell debris by centrifugation (17000 g x 30 min at 4°C), the supernatant was applied onto a Ni-NTA resin (Qiagen). After 10 CV washes with buffer A supplemented with 20 mM imidazole, the bound protein was eluted with buffer A supplemented with 200 mM imidazole. The eluted protein was concentrated and using using Amicon Ultra-15 centrifugal filters (3 kDa cut-off, Merck-Millipore) and then buffer was exchanged to buffer B (50 mM Tris-HCl pH 8.0. 5 mM CaCl_2_). The protein was loaded onto an anion-exchange column (MonoQ 10/100 GL, GE healthcare) equilibrated with buffer B, and eluted with a 0.1-0.4 M linear gradient of NaCl. The eluted protein was concentrated and treated with thrombin (Nacalai Tesque) at 4°C for overnight to cleave the His tag. The sample was further purified with a size exclusion column equilibrated with 20 mM Tris-HCl 150 mM and 5 mM CaCl_2._ The eluted protein was concentrated and stored at −80°C.

### SEC-MALS/SAXS

Size-exclusion chromatography coupled with multiangle light scattering (SEC-MALS) analysis was performed by using a high performance liquid chromatography (HPLC) system, Alliance 2695 (Waters) with DAWN HELEOS II (Wyatt Technology) and 2414 Refractive Index (RI) detector and 2489 UV/Visible detector (Waters). Samples (1.6∼1.7 mg/ml x 10 µl) wewe loaded on a Superdex 200 increase 3.2/300 column (Cytiva) equilibrated with SEC buffer (20 mM Tris-HCl pH 7.5, 150 mM NaCl, 10 mM CaCl_2_ and 0.02% GDN) under a flow rate at 0.1 ml/min. The conjugate analysis of the molecular masses by combining MALS-UV-RI signals was carried out by ASTRA 6.1 (Wyatt Technology) with a dn/dc value of 0.185 mL/g for protein and 0.17 mL/g for GDN, and an extinction coefficient (1 mg/ml) of 0.766 for protein and 0.01014 for GDN. Although the dn/dc value of GDN was arbitrarily used based on the other detergents, this value did not affect the calculation of the molecular masses of the protein.

Size-exclusion chromatography combined with small-angle X-ray scattering (SEC-SAXS) was performed on the BL-15A2 beamline at the photon factory, KEK (Tsukuba, Japan)^57^. The sample (1.6∼1.7 mg/ml x 50 µl) was loaded on a Superdex 200 increase 3.2/300 column equilibrated with the SEC buffer using a HPLC system Nexera-i (SHIMADZU) under the flow rate at 0.03 ml/min. A fiber spectrometer, QEpro (Ocean Insight) mounted at an angle of 45° to the sample cell was also utilized to obtain the concentration for each frame. The sample-to-detector distance was set to 2650 mm, and the measured X-ray wavelength was adjusted to 1.0 Å. The 2D scattering intensity images were recorded using PILATUS3 2M (DECTRIS) with an exposure time of 3 seconds. UV-visible absorption spectra were also recorded at 3-second intervals with a 1-second integration time. All scattering images were azimuthally averaged to convert the one-dimensional scattering intensity data, and then the subtraction of the background profile was also calculated. The absolute intensity calibration was performed using water as a standard. These processes were carried out using SAngler^58^. Scattering profiles above the half of the elution peaks were averaged by using MOLASS^59^ The radius of gyration (R_g_) and and the maximum molecular dimension (*D*_max_) from the Guinier approximation and the pair distribution function (*P*(r) were calculated by AUTORG and PRIMUSqt from ATSAS, respectively (Extended Data Fig. 1h)^60^.

### Colorimetric analysis

To detect the metal ions contained in the purified sample of MCFD2, PAR analysis was conducted^61^. The purified sample of MCFD2 (50 µM) was denatured with 4 M guanidine-HCl at pH 7.6 for 5 min at 37°C. The denatured sample was mixed with a freshly prepared solution of (4-(2-pyridylazo) resorcinol (PAR) (Tokyo Chemical industry) (final 100 µM) and then the UV-vis spectrum was immediately measured with a Hitachi UV3900. Note: Due to the relatively weak affinity, the amount of the bound metal ions (probably Zn^2+^ or Ni^2+^) in the purified MCFD2 samples varied between each purification.

### Isothermal titration calorimetry ITC

Isothermal titration calorimetry (ITC) experiments were carried out using the MicroCal™ iTC200 calorimeter (Malvern) at 293 K. After a 0.4 µL initial injection, 2 µL of ZnCl_2_ solution (500 µM) was injected into the EDTA-treated MCFD2 solution (15 µM) in 20 mM Tris–HCl pH 7.5, 150 mM NaCl and 0.2 mM CaCl_2_, at 180s intervals with stirring 750 rpm. The data analysis was performed with Microcal Origin Software (version 7.0) using a one-site binding model. The experiments were repeated at least twice with similar results.

### Cryo-EM sample preparation and data collection

The purified samples of full-length ERGIC-53 with or without MCFD2 (4∼5 mg/ml) were mixed with 3 mM Fluorinated Fos-Choline-8 (Anatrase), and then 3 µl aliquot of the sample was applied to glow-discharged QuantiFoil R1.2/1.3 Au 300 mesh grids. The grids were blotted for 4 s and immersed in liquid ethane using Vitrobot Mark IV systems (FEI/Thermo Fisher) operated at 4 °C and 100% humidity. The grids were initially screened using Talos Arctica (FEI) with a K2 direct electron detector (Gatan). The best grids were loaded to a Titan Krios (FEI) microscope operated at 300 kV and equipped with a Gatan Quantum-LS Energy Filter (GIF) and a Gatan K3 BioQuantum direct electron detector. Movies were automatically collected using SerialEM software^62^. For the sample of the ERGIC-53 1′H34-MCFD2 complex (1.7 mg/ml), two rounds of 3 µl aliquot loading and manual blotting were performed, and then a 3 µl aliquot of the sample was applied to glow-discharged QuantiFoil R1.2/1.3 Au 200 mesh grids, as previously reported ^63^. The grids were blotted for 5 s and immersed in liquid ethane using Vitrobot Mark IV systems (FEI/Thermo Fisher) operated at 4 °C and 100% humidity. Grid observation and data collection were performed with a CRYO ARM™ 300II (JEOL) operated at 300 kV and equipped with a JEOL in-column Omega energy filter and a Gatan K3 BioQuantum detector. Movies were automatically collected using SerialEM software^62^. The data collection parameters are summarized in Table 1.

**Table 1.** Cryo-EM data collection, refinement and validation statistics.

|  | Head region<br>(form A)<br>(EMDB-36467)<br>(PDB 8JP4) | Head region (form<br>B)<br>(EMDB-36467)<br>(PDB 8JP5) | Head region<br>(substate A)<br>(EMDB-36468)<br>(PDB 8JP6) | Head region<br>(substate B)<br>(EMDB-36469)<br>(PDB 8JP7) |
| --- | --- | --- | --- | --- |
| **Data collection and processing** |  |  |  |  |
| Magnification | 105,000 | 105,000 | 105,000 | 105,000 |
| Voltage (kV) | 300 | 300 | 300 | 300 |
| Electron exposure (e-/Å<sup>2</sup>) | 47.4 | 47.4 | 47.4 | 47.4 |
| Defocus range (μm) | -0.8~ -1.6 | -0.8~ -1.6 | -0.8~ -1.6 | -0.8~ -1.6 |
| Pixel size (Å) | 0.83 | 0.83 | 0.83 | 0.83 |
| Symmetry imposed | C2 | C2 | C1 | C1 |
| Initial particle images (no.) | 2,132,031 | 2,132,031 | 2,132,031 | 2,132,031 |
| Final particle images (no.) | 89361 | 81444 | 9734 | 9783 |
| Map resolution (Å) | 2.53 | 2.59 | 3.29 | 3.51 |
| FSC threshold (0.143) |  |  |  |  |
| Map resolution range (Å) | 2.4~4.8 | 2.4~4.8 |  |  |
| **Refinement** |  |  |  |  |
| Initial model used (PDB code) | 3WNX |  |  |  |
| Model resolution (Å) | 3.0 | 3.2 | 3.3 | 3.5 |
| FSC threshold (0.5) |  |  |  |  |
| Model resolution range (Å) |  |  |  |  |
| Map sharpening *B* factor (Å<sup>2</sup>) | -68.9 | -73 | -65 | -55 |
| Model composition |  |  |  |  |
| Non-hydrogen atoms | 13500 | 13490 | 13500 | 13476 |
| Protein residues | 1678 | 1676 | 1678 | 1674 |
| Ligands | 20 | 20 | 20 | 20 |
| *B* factors (Å<sup>2</sup>) |  |  |  |  |
| Protein | 185.8 | 185.8 | 180.4 | 254.5 |
| Ligand | 132.31 | 132.8 | 144 | 202.9 |
| R.m.s. deviations |  |  |  |  |
| Bond lengths (Å) | 0.002 | 0.003 | 0.003 | 0.005 |
| Bond angles (°) | 0.582 | 0.577 | 0.593 | 0.636 |
| Validation |  |  |  |  |
| MolProbity score | 1.53 | 1.77 | 1.48 | 1.61 |
| Clashscore | 5.4 | 5.03 | 4.45 | 5.64 |
| Poor rotamers (%) | 1.8 | 2.35 | 0.14 | 0.62 |
| Ramachandran plot |  |  |  |  |
| Favored (%) | 97.8 | 96.6 | 96.2 | 95.7 |
| Allowed (%) | 2.2 | 3.4 | 3.8 | 4.3 |
| Disallowed (%) | 0.0 | 0.0 | 0.0 | 0.0 |

| | Head region<br>(substate C)<br>(EMDB-36471)<br>(PDB 8JP8) | Head region<br>(Substate D)<br>(EMDB-36472)<br>(PDB (8JP9) | Full length<br>(EMDB-36479)<br>(PDB 8JPG) | ΔH34 mutant<br>(EMDB-36482) |
| --- | --- | --- | --- | --- |
| **Data collection and processing** |  |  |  |  |
| Magnification | 105,000 | 105,000 | 105,000 | 60,000 |
| Voltage (kV) | 300 | 300 | 300 | 300 |
| Electron exposure (e<sup>-</sup>/Å<sup>2</sup>) | 47.4 | 47.4 | 47.4 | 50 |
| Defocus range (μm) | -0.8~ -1.6 | -0.8~ -1.6 | -0.8~ -1.6 | -0.8 - -1.6 |
| Pixel size (Å) | 0.83 | 0.83 | 0.83 | 0.788 |
| Symmetry imposed | C1 | C1 | C1 | C1 |
| Initial particle images (no.) | 2,132,031 | 2,132,031 | 168,458 | 1,279,058 |
| Final particle images (no.) | 10,713 | 13,895 | 29762 | 138,608 |
| Map resolution (Å) | 3.39 | 3.37 | 6.72 | 3.53 |
| FSC threshold (0.143) |  |  |  |  |
| Map resolution range (Å) |  |  |  |  |
| **Refinement** |  |  |  |  |
| Initial model used (PDB code) |  |  |  |  |
| Model resolution (Å) | 3.4 | 3.4 | 8.9 |  |
| FSC threshold (0.5) |  |  |  |  |
| Model resolution range (Å) | -54 | -56 |  |  |
| Map sharpening *B* factor (Å<sup>2</sup>) |  |  |  |  |
| Model composition |  |  |  |  |
| Non-hydrogen atoms | 13500 | 13500 | 18068 |  |
| Protein residues | 1678 | 1678 | 2252 |  |
| Ligands | 20 | 20 | 20 |  |
| *B* factors (Å<sup>2</sup>) |  |  |  |  |
| Protein | 214.1 | 218.8 | 1179.6 |  |
| Ligand | 221.1 | 185.6 | 1124.6 |  |
| R.m.s. deviations |  |  |  |  |
| Bond lengths (Å) | 0.002 | 0.03 | 0.004 |  |
| Bond angles (°) | 0.516 | 0.507 | 0.781 |  |
| Validation |  |  |  |  |
| MolProbity score | 1.2 | 1.19 | 1.87 |  |
| Clashscore | 2.78 | 2.44 | 11.0 |  |
| Poor rotamers (%) | 0.07 | 0.14 | 0.1 |  |
| Ramachandran plot |  |  |  |  |
| Favored (%) | 97.3 | 97.1 | 95.6 |  |
| Allowed (%) | 2.7 | 2.9 | 4.4 |  |
| Disallowed (%) | 0.0 | 0.0 | 0.0 |  |

### EM image processing

Movies were aligned using beam-induced motion correction implemented in RELION v3.1 and 4.0^64,65^ and contrast transfer function parameters were estimated using Patch-based CTF estimation in cryoSPARC^66^. The following image processing was mainly performed by using cryoSPARC. Bayesian polishing was performed by RELION v3.1 and 4.0 with the help of UCSF pyem tool^67^ for data conversion from cryoSPARC to Relion.

For the sample of ERGIC-53 in complex with MCFD2, 6345 movies were collected. For analysis of the head and TM regions, blob-based autopicking and subsequent 2D classification were performed by using 800 movies to create templates for template-based autopicking in cryoSPARC. A total of 2,132,031 particles were automatically picked by the template picker in cryoSPARC and extracted at a pixel size of 3.32 Å followed by 2D classification. Good classes with clear 2D average images of the head region were selected and subjected to ab-initio modeling and subsequent heterogeneous refinements. Particles belonging to forms A and B were re-extracted at a pixel size of 0.996 Å and subjected to non-uniform (NU) refinement^68^. The refined particles were subjected to Bayesian polishing and then NU refinement with global CTF and particle defocus refinement. Consequently, the final maps of forms A and B were refined to 2.53 and 2.59 Å resolution, respectively. The maps were sharpened and locally filtered based on the local resolution in cryoSPRAC.

For its full-length particle analysis, full-length ERGIC-53 particles were manually picked from 50 micrographs. Particles showing full-length-like structures were selected from 2D classification and subjected to Topaz training^40^. A trained model of Topaz was further trained by repeating auto-picking by Topaz and 2D classification using 50, 100, 600 micrographs. Subsequently, full-length particles were picked by Topaz from all micrographs, and extracted at a pixel size of 2.49 Å. 2D classification generated clear 2D average images of full-length ERGIC-53 particles. Particles belonging to the straight conformation were selected and subjected to homogeneous refinement and local refinement. The final map was refined to 6.8 Å resolution.

For the ERGIC-53ΔH34-MCFD2 complex, 6175 movies were collected and CTF parameters were estimated with CTFFIND4. Particles were initially picked using Laplacian-of-Gaussian-(LoG) based auto-picking in RELION 4 to create 2D templates. A total of 127,9058 particles were picked and extracted at a pixel size of 3.142 Å and subjected to 2D classification. Good 2D classes were selected and subjected to ab-initio modeling and subsequent heterogeneous refinements. The best particles were re-extracted at a pixel size of 1.3791 Å and subjected to homogenous refinement. The refined particles were subjected to Bayesian polishing, CTF refinement, defocus refinement and homogenous refinement. The final map (consensus map) was reconstructed at 3.51 Å resolution.

To gain insight into the structural viability of all the obtained EM maps, the refined particles were subjected to 3D variability analysis in cryoSPRAC^42^. In the head region analysis, particles were further classified by cluster analysis in 3DVA (12 classes), resulting in the determination of four substate structures at 3.3∼3.4 Å resolution.

### Model building, refinement and visualization

For the CRD region and MCFD2, the previously published crystal structure of the CRD and MCFD2 complex (PDB ID 3WNX) was docked in the EM map. The S-H1 and S-H2 regions were manually built by using Coot^69^. The models were refined with Servalcat^70^ and phenix.real_space_refine^71^ and further manually corrected with Coot. For the building of the full-length model, partial tetrameric models of S-H3-SH4, and S-H4-TM were predicted by using Colabfold with AF2 multimer ^72^. Then partial models were assembled and manually docked in the EM map. The model was further refined with phenix_real_space refine as rigid bodies. Structural figures were prepared with PyMOL (http://www.pymol.org) and Chimera X ^73^. Sequence alignment was performed and visualized with Jalview^74^

### Secretion assay

HEK293T WT or ERGIC-53 KO cells (6 x 10^5^ cells) were plated on 6 well plates and transfected with 100 ng of the plasmid expressing FV-HiBIT /AAT-HiBiT and 200 ng of the plasmid expressing ERGIC-53 WT or each of its mutant. After 24 h incubation, the medium was removed, and cells were washed with Expi293 medium and incubated in 1 ml of Expi293 medium for 4h. In the Zn^2+^ supplement conditions, 10 µM ZnCl_2_ was added in the medium. After harvest of the conditioned mediums (CMs), the remained cells were washed with PBS and lysed in a lysis buffer (PBS, 1% Triton (Thermo Fisher) and protease inhibitor cocktail (Nakarai Tesque), and then briefly sonicated. The total volume of the cell lysates was adjusted to the same volume with the CMs. The amounts of secreted and intracellular HiBiT-tagged FV/AAT proteins were quantified by using the Nano-Glo HiBiT Extracellular Detection System or Nano-Glo HiBiT lytic assay system, respectively. Luminescence from HiBiT-tagged proteins was measured with GloMax Navigator (Promega). Luminescence from CMs and lysates of nontransfected cells were used for background correction.

Secreted and intracellular HiBiT-tagged FV or AAT proteins as well as expressed ERGIC-53 were also detected by Western blotting with the Nano-Glo® HiBiT Blotting System (LgBiT protein) and anti-Factor V (Abcam, ab108614, 1:3300) or anti-LMAN1 (abcam, ab125006, 1:5000) antibodies. Total proteins were detected by stain-free technology (Bio-Rad). The experiments were performed at biologically independently at least three time. Statistical analysis was performed with Graphpad prism 9.3, using a two-tailed unpaired t-test for comparison of two groups or one-way ANOVA followed by Dunnett’s test for the other comparisons.

### Data and materials availability

The cryo-EM density maps have been deposited in the Electron Microscopy Data Bank (EMDB) under accession codes EMD-36467 (form A), EMD-36468 (form B), EMD-36469 (substate A), EMD-36470 (substate B), EMD-36471 (substate C), EMD-36472 (substate D), EMD-36479 (full length) and EMD-36482 (the ΔH34 mutant). The atomic coordinates have been deposited in the Protein Data Bank under accessions 8JP4 (form A), 8JP5 (form B), 8JP6 (substate A), 8JP7 (substate B), 8JP8 (substate C), 8JP9 (substate D), and 8JPG (full length). Raw movies have been deposited in EMPIAR-11645 (the ΔH34 mutant) and EMPIAR-11646 (full-length ERGIC-53).

## Acknowledgement

We would like to thank staff scientists at the University of Tokyo’s cryo-EM facility, especially, A. Tsutsumi, K, Kobayashi, H. Yanagisawa, M. Kikkawa and R. Danev; T. Yokoyama, K. Nanatani, J. Inoue, S. Koshiba, and M. Yamamoto for management of the cryo-EM facility at Advanced Research Center for Innovations in Next-Generation Medicine in Tohoku University; and Y. Amagai and M. Michio for help with cell cultures. This work was supported by Grants-in-Aid for Scientific Research (JP18K06075 to S.W.). Grant-in-Aid for Transformative Research Areas (JP21H05253 to I.K. and S.W.) from the MEXT of Japan, a research grant from the Naito foundation (to S.W.), a research grant from Mochida memorial foundation for medical and pharmaceutical research (to S.W), and Platform Project for Supporting Drug Discovery and Life Science Research (Basis for Supporting Innovative Drug Discovery and Life Science Research (BINDS)) from AMED under grant number JP21am0101115 (support number 1025) JP21am0101071 (support number 2078), and JP21am0101095.

## Author contributions

S.W. designed the research and performed almost all experiments, cryo-EM data collection at Tohoku university, image processing and model building and refinement. Y.K. performed grid preparation/optimization and cryo-EM data collection at University of Tokyo under the supervision of O.N.. K.Y performed SEC-MALS/SAXS analysis at KEK under the supervision of N.S. M.I performed plasmid construction and secretion assays. S.W. and K.I. wrote the manuscript with the help from all other authors. S.W and K.I supervised this work.

## Competing interests

We declare that there are no competing interests related to this work.

**Correspondence and requests for materials** should be addressed to Satoshi Watanabe or Kenji Inaba

## Extended Data figure legends

**Extended Data Fig. 1.**
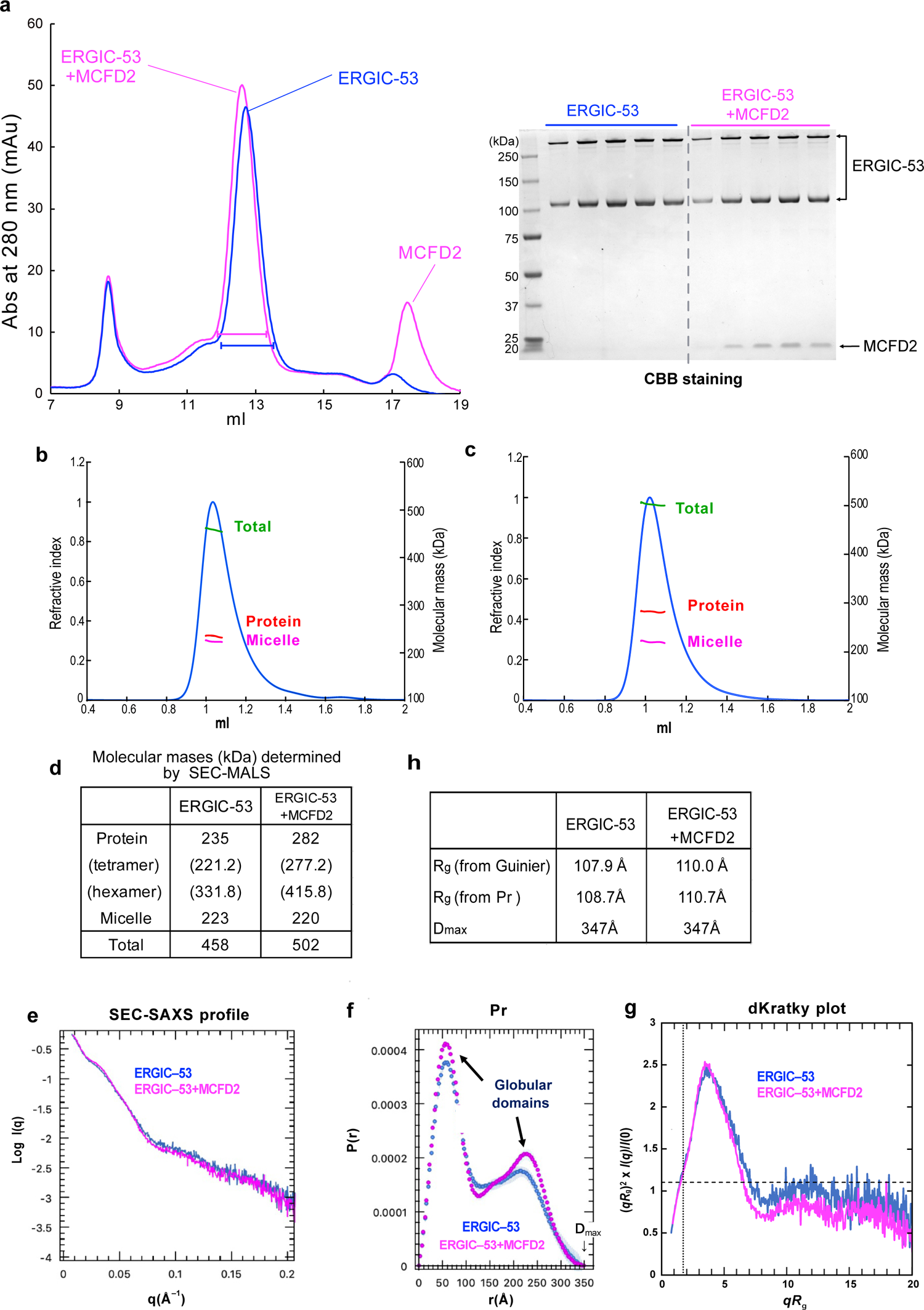
SEC-MALS/SAXS analysis of recombinant human ERGIC-53 in complex with MCFD a) SEC profiles of purified ERGIC-53 (blue) and the mixture of ERGIC-53 and MCFD2 (pink). The right panel shows SDS-PAGE analysis of each peak fraction, detected by CBB staining. b,c) The SEC-MALS profile of ERGIC-53 (a) and its complex with MCFD2(b). Masses of the peak fractions determined by SEC-MALS analysis are indicated by the green line for total mass, red for protein mass and magenta line for micelle, respectively. d) Summary of the determined molecular masses. e) Experimental SAXS profiles of ERGIC-53 (blue) and ERGIC-53 in complex with MCFD2 (magenta). f) Pair distance distribution functions P(r) of ERGIC-53 (blue) and the complex (magenta) determined from the SAXS profiles. g) The dimensionless Kratky plots of ERGIC-53 (blue) and the complex (magenta) determined from the SAXS profiles. The dotted and dashed lines represent the positions of √3 and 3/e, respectively, and the peak appears at this cross point for a typical globular protein. h) Summary of radius of gyration (R_g_) and maximum dimension D_max_.

**Extended Data Fig. 2.**
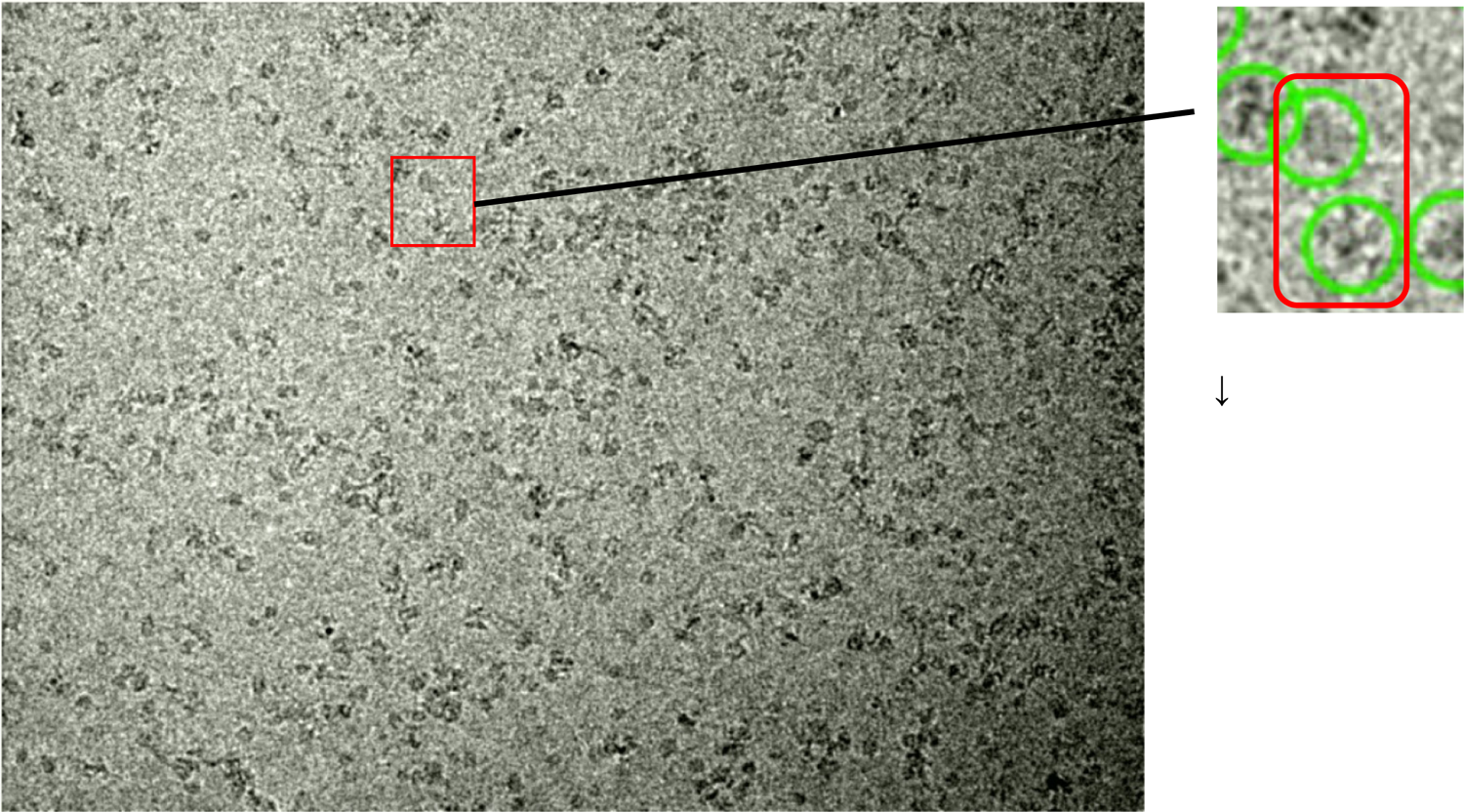
Cryo-EM analysis of ERGIC-53 in complex with MCFD2 A representative motion-corrected micrograph of ERGIC-53 with MCFD2. The inset shows a close-up view of a dumbbell-like particle of full-length ERGIC-53, indicated by a red square. Green circles represent particles picked by blob-based autopicking.

**Extended Data Fig. 3.**
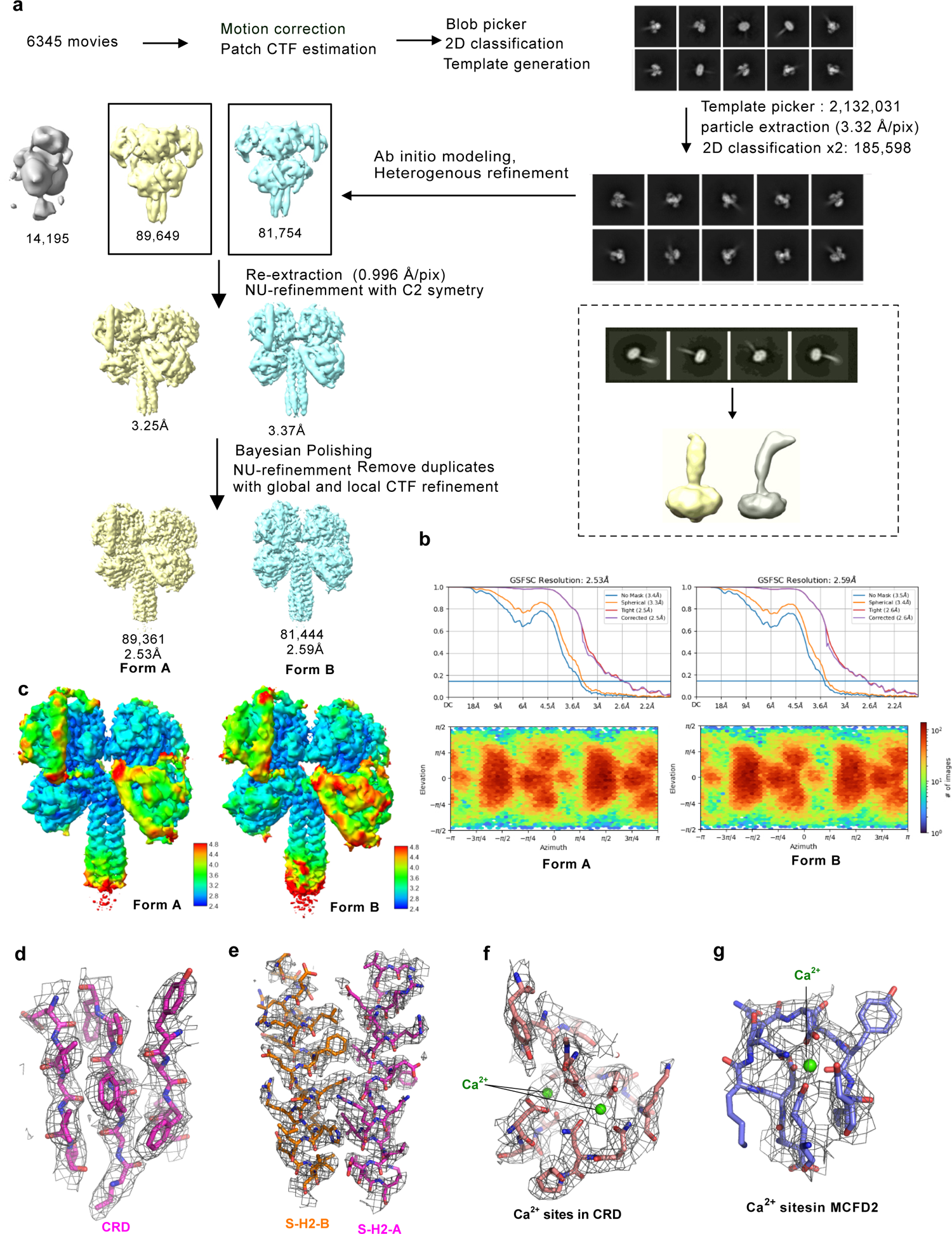
Cryo-EM image processing of the head region of ERGIC-53 in complex with MCFD2 a) Workflow of the image processing of each globular region. Representative 2D class average images and refined maps at each step are shown. b) Gold standard Fourier shell correlation (GSFSC) resolution plots (upper panels) and Euler angle distributions (lower panels) of the final maps of form A and B, calculated with cryoSPARC. c) Local resolution of the final maps calculated with the local resolution tool in cryoSPARC. d∼g) Representative EM density maps and models around (d) the center of CRD, (e) the central four-helix coiled coil, (f) Ca^2+^ binding site in the CRD, and (g) Ca binding site in MCFD2

**Extended Data Fig. 4.**
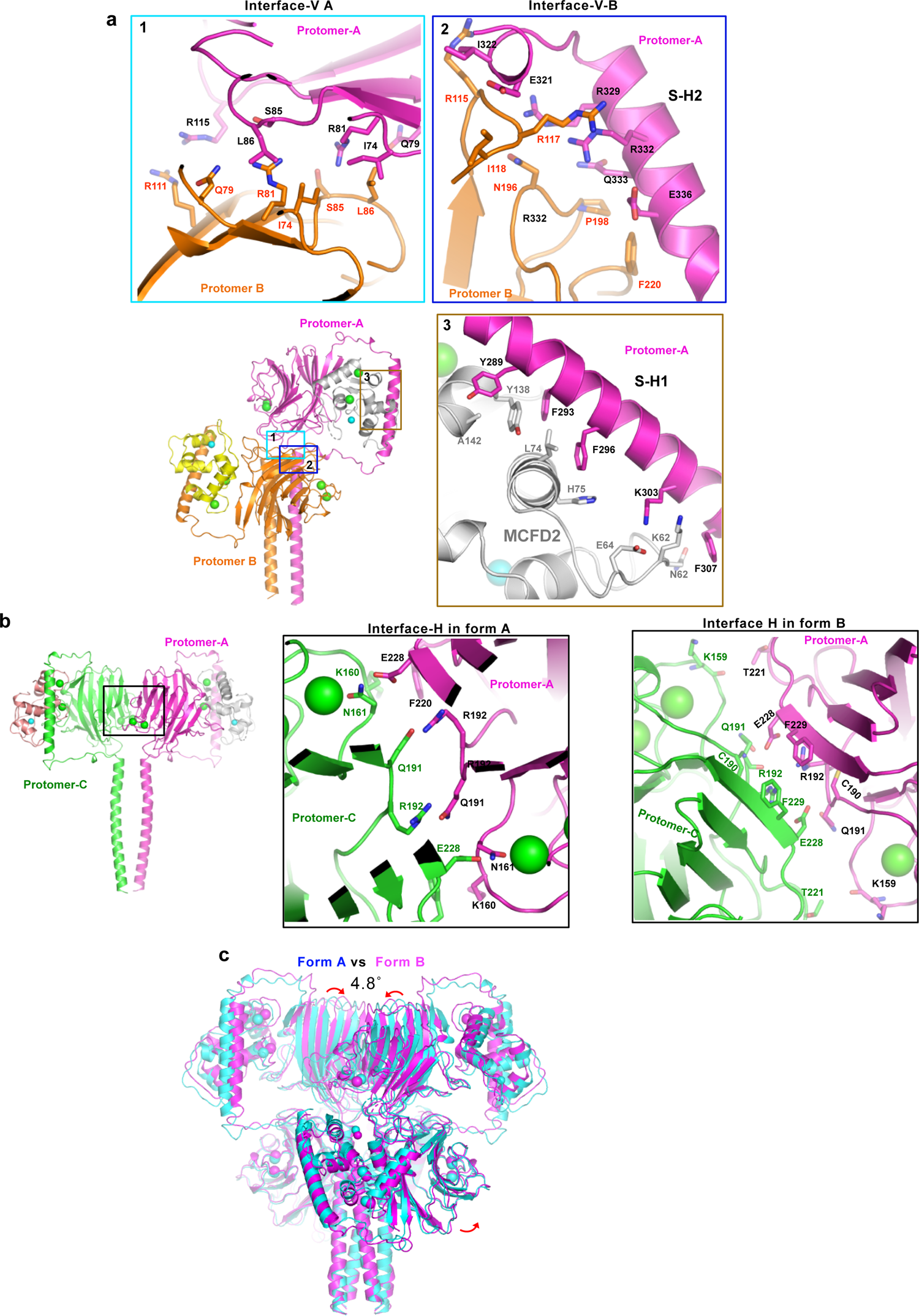
Details of domain interfaces a) Details of the vertical interface (interface-V) and SH-1-MCFD2 interaction. The left lower panel shows the overall structure of the head region dimer. The other three panels show close-up views at the cyan, blue, and yellow boxes, respectively; 1: interface between the upper and lower CRD; 2: interface between the lower CRD and S-H2 helix; 3: interaction between S-H1 and MCFD2. Residues involved in the interface are shown in stick models. b) Details of the horizontal interface (interface-H). (left) The head region dimer formed by interface-H between the upper CRDs in form A; (middle and right) close-up view a of the interface H at the black box in form A (left) or form B (right). Residues involved in the interface are shown in stick models. Bound calcium ions are represented by green spheres. c) Comparison of the form A (cyan) and form B (magenta) structures, in which their the central-coiled coils are superimposed on each other.

**Extended Data Fig. 5.**
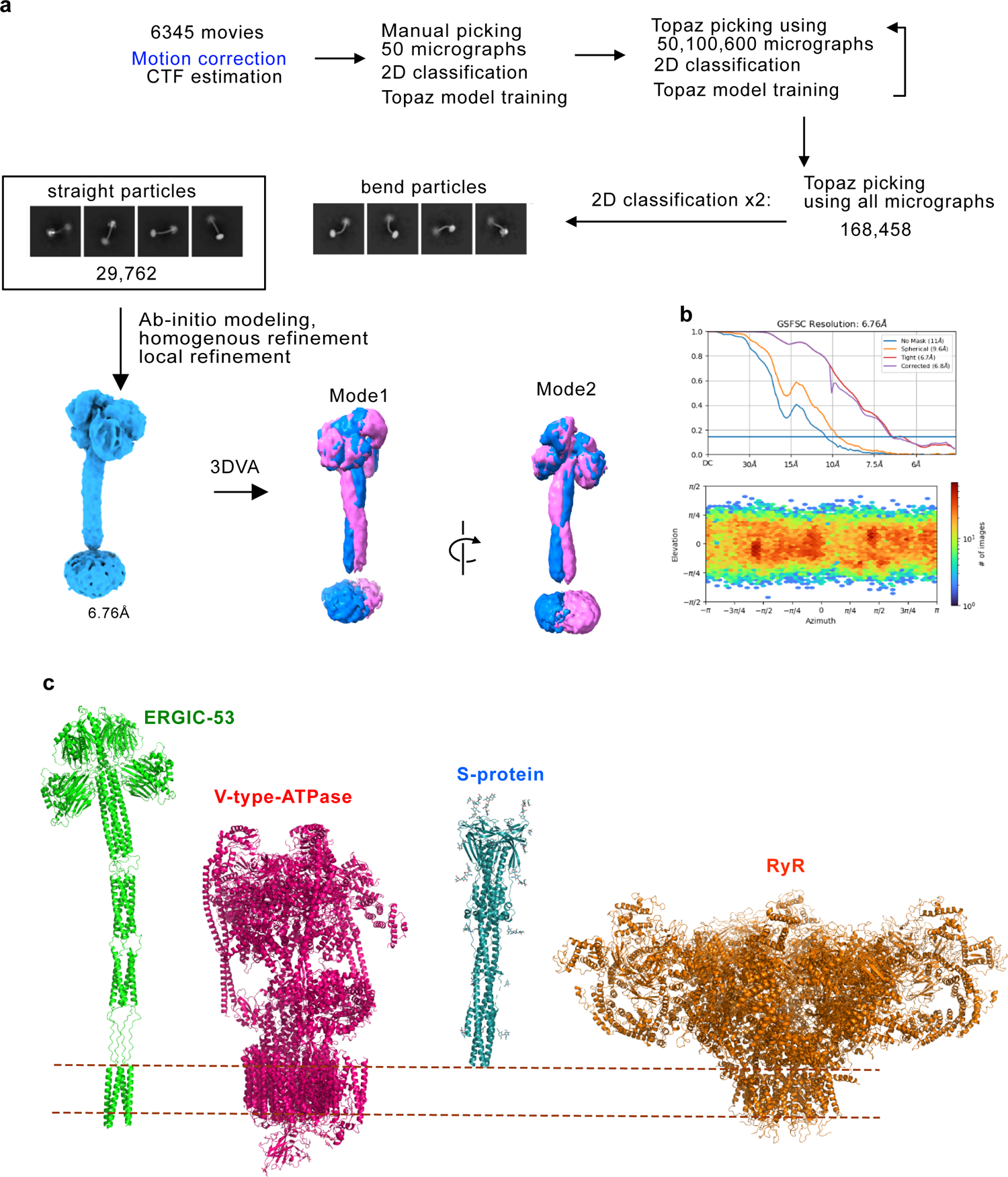
Imaging processing of the full-length particles of ERGIC-53. a) Workflow of the image processing of the full-length ERGIC-53 particles. Representative 2D class average images and refined maps at each step are shown. b) GSFSC resolution plots (upper panels) and Euler angle distributions (lower panels) of the final map at 6.7Å resolution. c) Comparison of the molecular height of full length ERGIC-53 complexed with MCFD2 (this study) with those of V-type ATPase (PDB: 6WM2), S-protein from SARS-COV-2 (PDB: 6XRA), and RyR (PDB: 5TB4).

**Extended Data Fig. 6.**
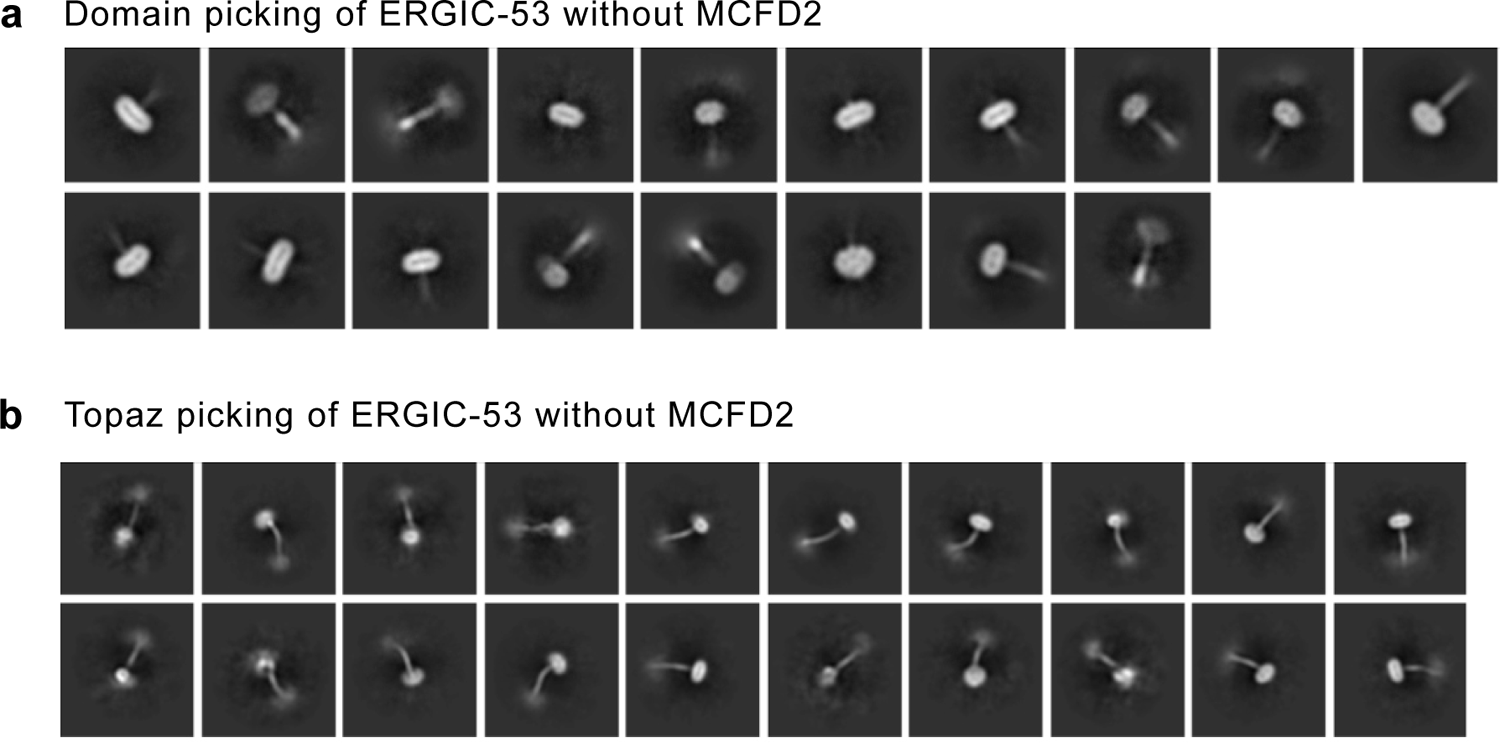
Cryo-EM SPA analysis of ERGIC-53 without MCFD2. Representative 2D class average images of the half regions of the particle(upper) and the full-length particles (lower).

**Extended Data Fig. 7.**
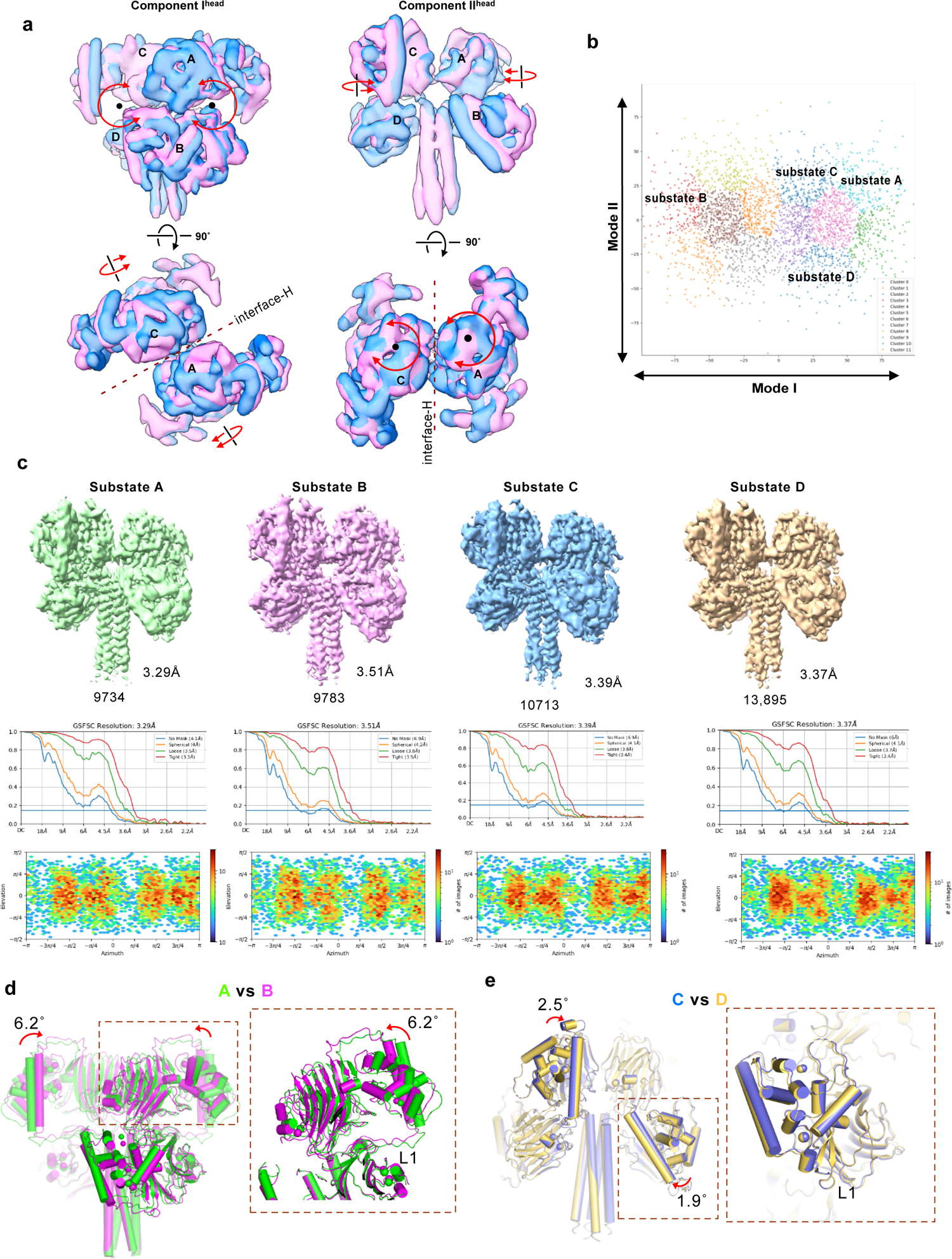
3DVA of the head region a) Results of 3DVA of the head region with two variability components I and II. The EM maps of the first (pink) and last frames (blue) of continuous conformational changes generated by 3DVA are displayed. arrows represent the rigid body rotation of each unit. b) Each dot represents a particle plotted according to the 3DVA clustering. c) Reconstruction and refinement of four substate structures of the head region. Particles belonging to each group (A, B, C, D) are indicated in the cluster plots in a. GSFSC resolution plots and Euler angle distributions of each substate structure are provided in the top, middle and lower panels, respectively. d) Comparison between substates A and B. e) Comparison between substates C and D.

**Extended Data Fig. 8.**
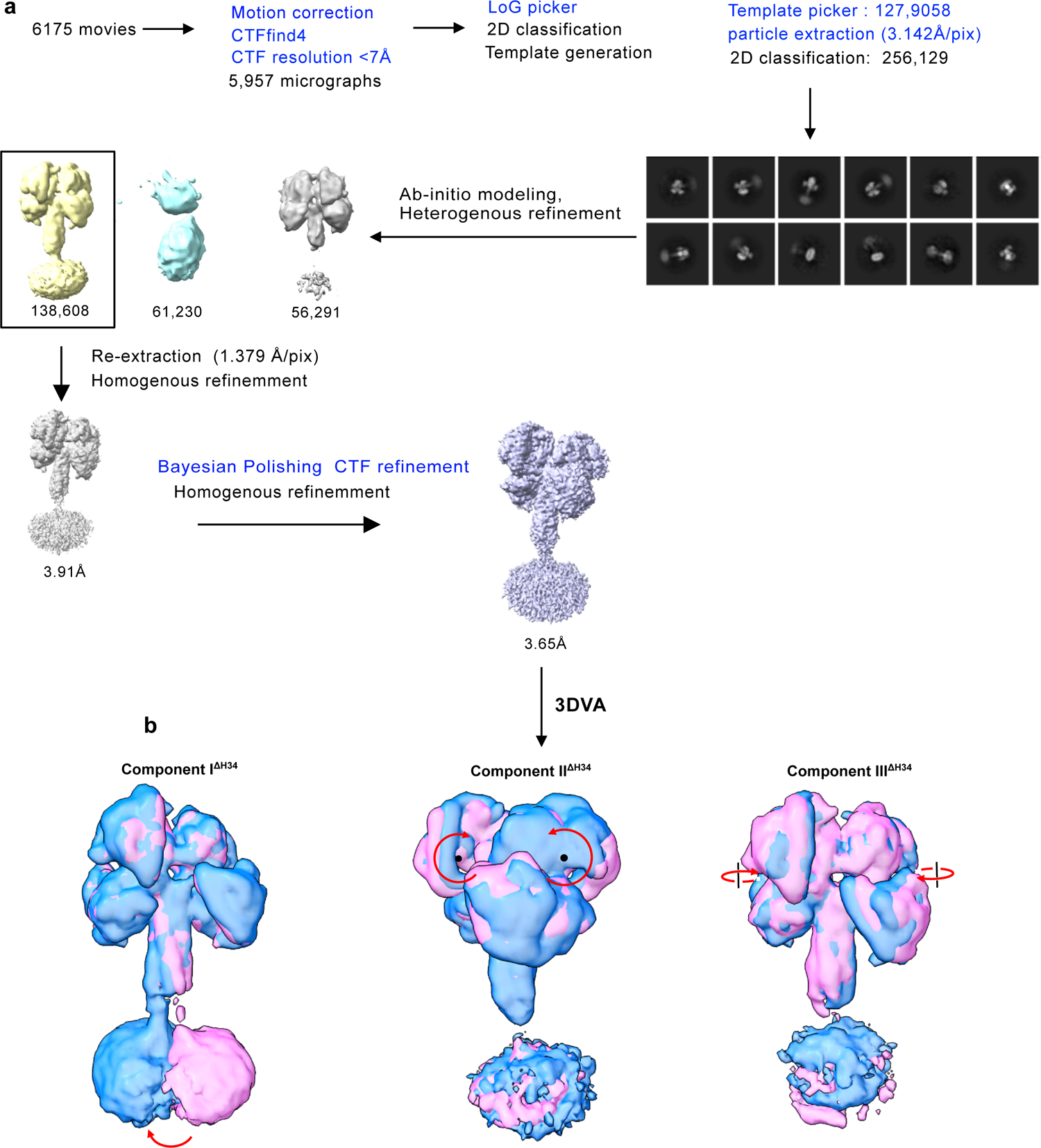
Cryo-EM image processing of the ERGIC-53 ΔH34 mutant with MCFD2 a) Workflow of the image processing of the ΔH34 mutant or ERGIC-53. Representative refined maps at each step are shown. b) The results of 3DVA of this mutant with three variable components I, II and III. The EM maps of the first (pink) and last frames (blue) of continuous conformational changes generated by 3DVA are displayed. Arrows represent rigid body rotation of each unit.

**Extended Data Fig. 9.**
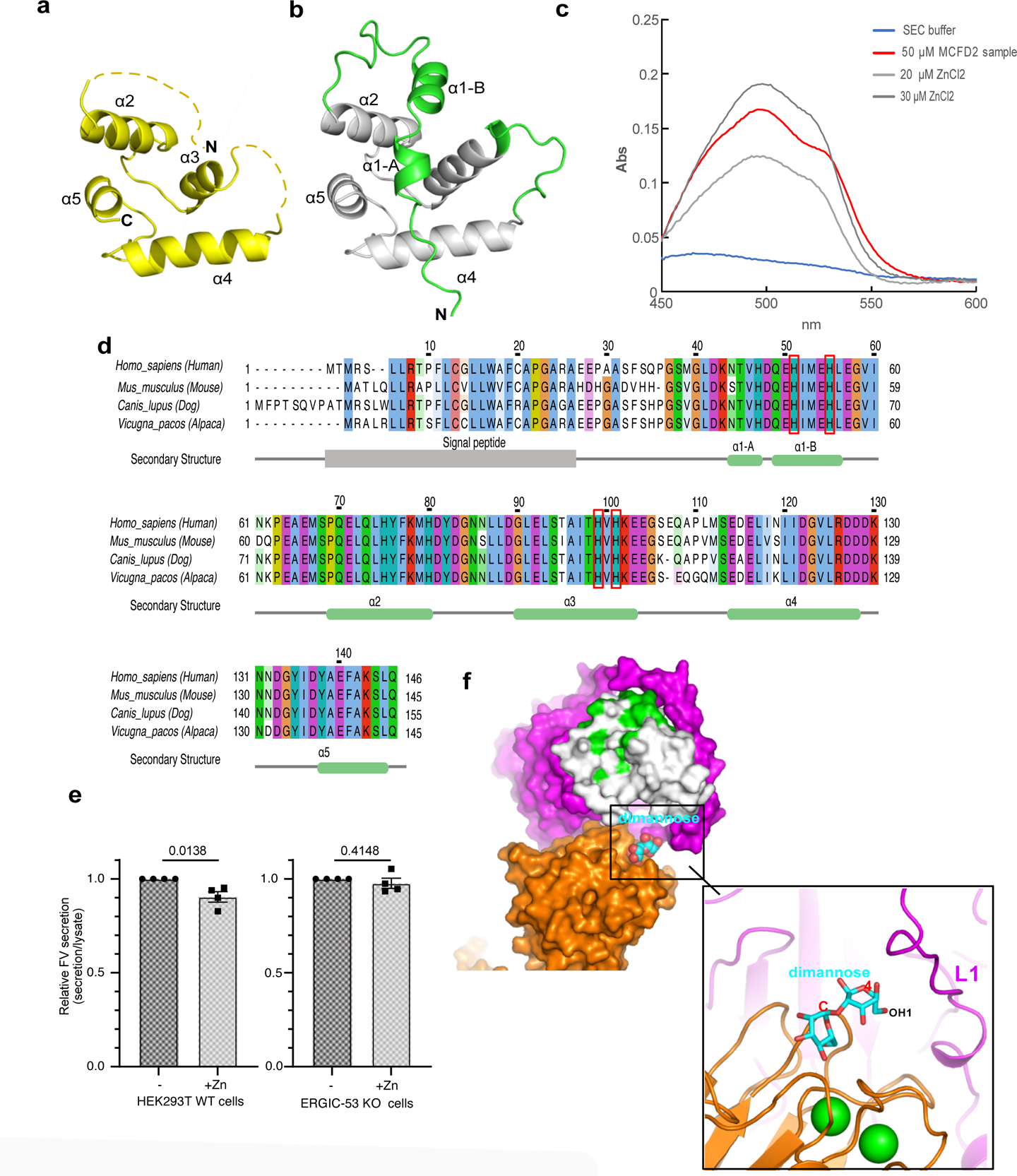
Details of Zn^2+^-dependent regulation of MCFD2 a) The original crystal structure of MCFD2 (PDB: 4YGE) b) A predicted structure of MCFD by Alphafold2. The residues that are missing in the original structures are shown in green. c) UV-vis spectrum of the PAR assay of purified denatured MCFD2 (red) or buffer (blue). d) Sequence alignment of MCFD2 homologues by Jalview. The four conserved Histidine residues are highlighted by red boxes. e) Secretion assay of HiBiT-tagged FV in HEK293T cells or ERGIC-53 KO with or without zinc supplementation. The amounts of secreted FV relative to that of intracellular FV under Zn supplementation conditions were quantified with a commercial HiBiT-tagged protein detection reagent, and normalized to the control conditions. Dots represent the individual data points. Error represents the standard error of the mean (N=4 biological replicates). An unpaired two-tailed t-test was used for statistical analysis. f) A docking model of dimannose in the present structure. The bound di-mannose is docked based on superposition of the crystal structure of the CRD with di-mannose (PDB: 4GKX). An inset shows a close-up view of di-mannose binding on the CRD, suggesting that the putative Man(4) moiety of Man_9_(GlcNAc)_2_ glycan lies near the L1 loop between the S-H1 and S-H2 helices.

**Extended Data Fig.10.**
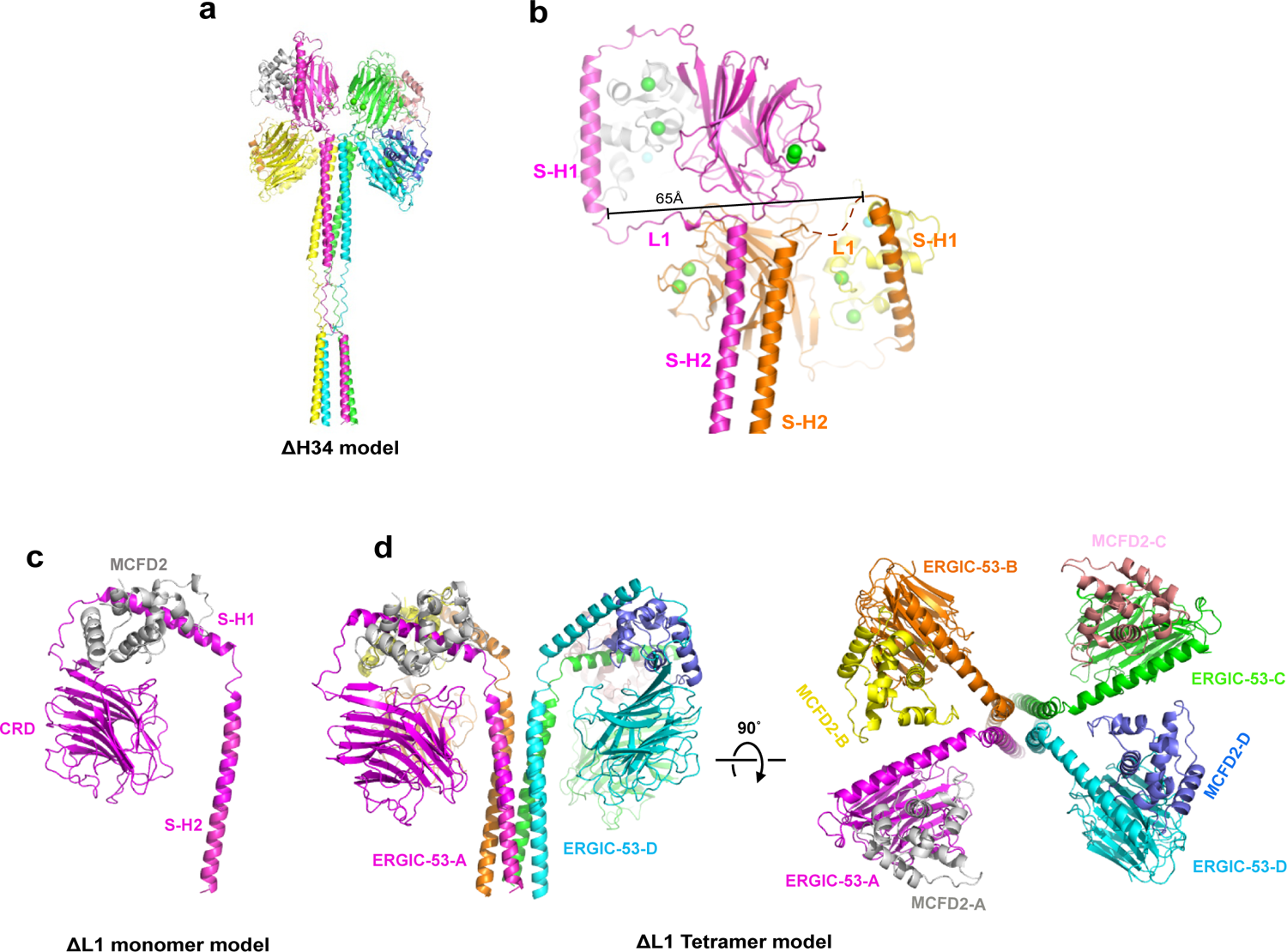
Structure models of truncated mutant of ERGIC-53 a) Overall structure model of the ERGIC-53 ΔH34 mutant. b) Close-up view of the L1 loop between two stalk helices (S-H1 and S-H2). In the present tetramer, the distance between the C-terminal ends S-H1 is ∼65Å, apart by the L1 loop. c) A model of the head region (CRD, SH1 and SH2) of the ΔL1 mutant tetramer with MCFD2 predicted by Colabfold v.1.5

**Extended Data Fig. 11.**
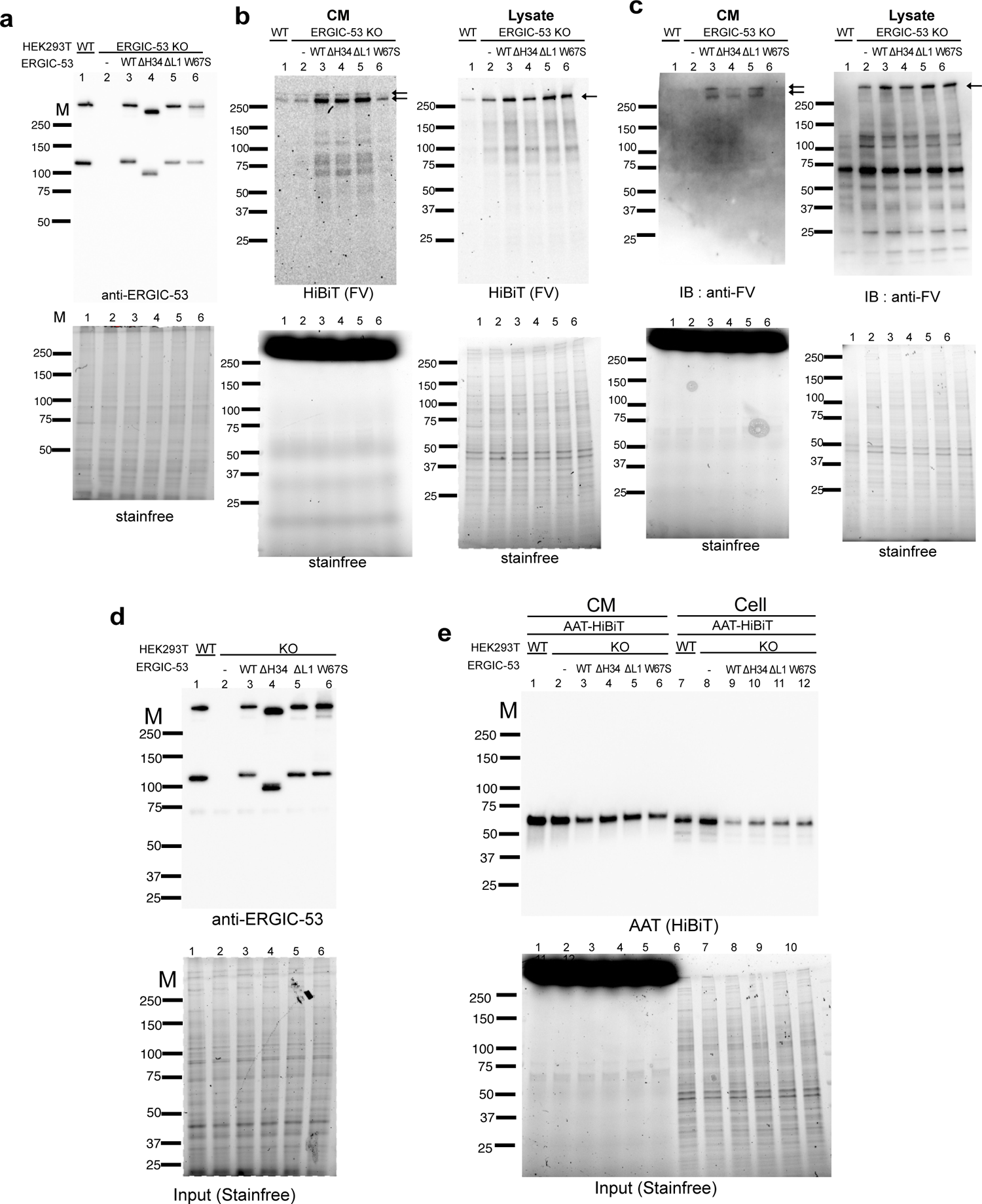
Rescue experiments of the secretion of FV-HiBiT and AAT-HiBiT a,b,c) Comparison of ecpression level of ERGIC-53 (a) and secreted and expressed FV (b). HEK293T cells (Abcam) transfected with FV-HiBiT and empty vector, and ERGIC-53 KO 293T cells transfected with FV-HiBiT and ERGIC-53, indicated mutant or empty vector were incubated in the serum-free Expi293 medium for 4 h. Conditioned media and cell lysates were resolved by nonreducing SDS‒PAGE (a) or reducing SDS‒PAGE (b, c) and analyzed with immunoblotting with anti-ERGIC-53 antibody (a), LiBiT proteins for HiBiT tag (b) or anti-FV antibody (c). Total protein in each lane was detected by stain-free technology. Note: The transfection efficiency of FV in HEK293T WT cells (Abcam) were significantly lower than that in the KO cells. Although the used anti-FV antibody failed to detect low levels of FV from these cells, HiBiT-blotting system showed higher sensitivity and detected secreted/expressed FV-HiBiT proteins in all lanes. d, e) Comparison of the expression level of ERGIC-53 (d) and secreted and expressed AAT-HiBiT (e). HEK293T cells transfected with AAT-HiBiT and empty vector, and ERGIC-53 KO 293T cells transfected with AAT-HiBiT and ERGIC-53, indicated mutant or empty vector were incubated in the serum-free Expi293 medium for 4h. Conditioned media and cell lysates were resolved by non-reducing (d) or reducing (e) SDS-PAGE and analysed with immunoblotting with anti-ERGIC-53 antibody and LgBiT proteins for HiBiT-tagged proteins.

**Extended Data Fig.12.**
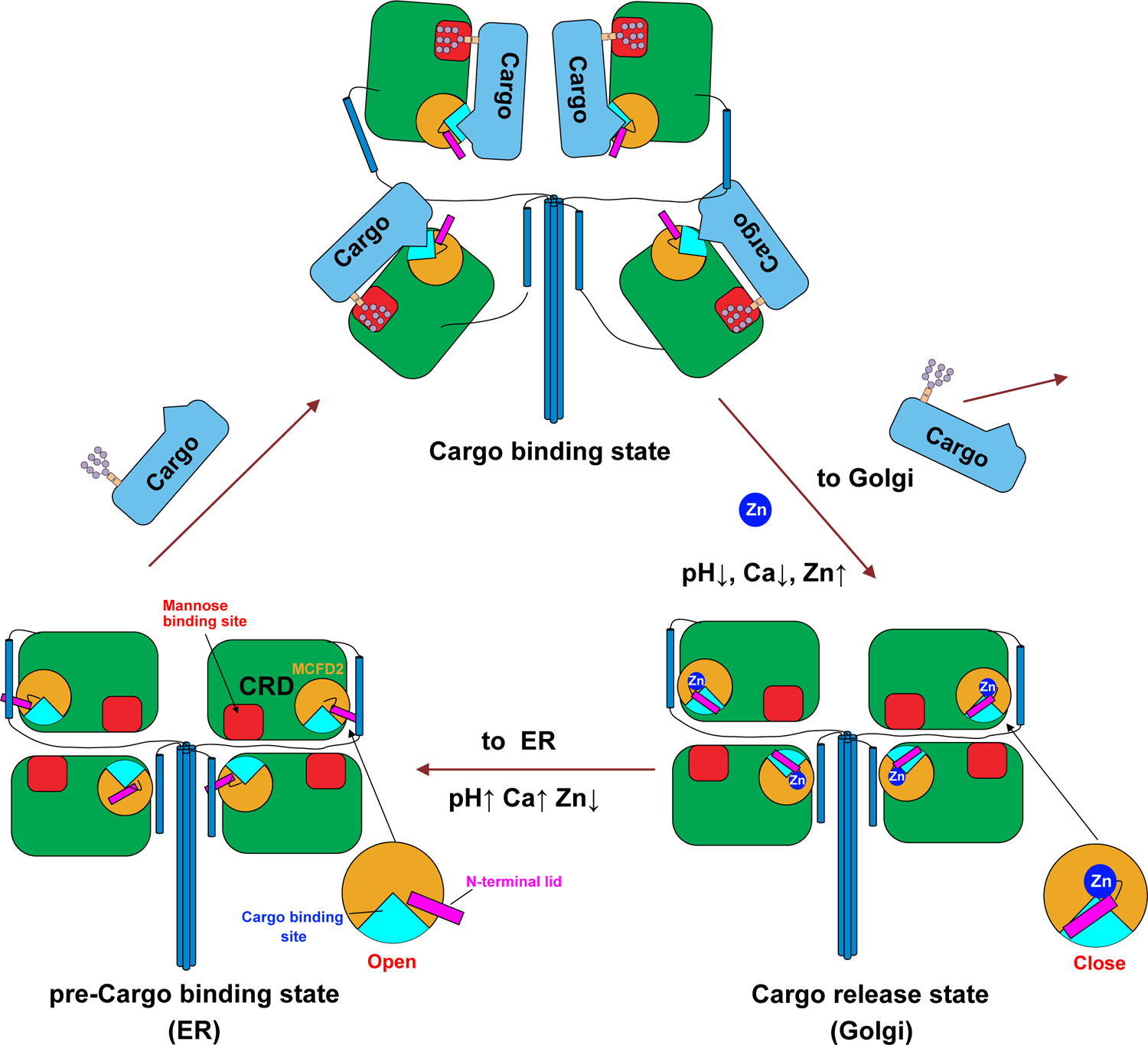
A working model of Zn^2+^, Ca^2+^ and pH-dependent regulation of cargo capture and release by the ERGIC-53 and MCFD2 complex (see text).

## References

1. Barlowe, C. & Helenius, A. Cargo Capture and Bulk Flow in the Early Secretory Pathway. Annu. Rev. Cell Dev. Biol. 32, 197–222 (2016).

2. Tang, V. T. & Ginsburg, D. Cargo selection in endoplasmic reticulum–to– Golgi transport and relevant diseases. J. Clin. Invest. 133, e163838 (2023).

3. Barlowe, C. K. & Miller, E. A. Secretory protein biogenesis and traffic in the early secretory pathway. Genetics 193, 383–410 (2013).

4. Okumura, M., Kadokura, H. & Inaba, K. Structures and functions of protein disulfide isomerase family members involved in proteostasis in the endoplasmic reticulum. Free Radic. Biol. Med. 83, 314–322 (2015).

5. Zhang, Y. C., Zhou, Y., Yang, C. Z. & Xiong, D. S. A review of ERGIC-53: its structure, functions, regulation and relations with diseases. Histol. Histopathol. 24, 1193–1204 (2009).

6. Zheng, C. & Zhang, B. Combined Deficiency of Coagulation Factors V and VIII: An Update. Semin. Thromb. Hemost. 39, 613–620 (2013).

7. Nichols, W. C. et al. Mutations in the ER–Golgi Intermediate Compartment Protein ERGIC-53 Cause Combined Deficiency of Coagulation Factors V and VIII. Cell 93, 61–70 (1998).

8. Nyfeler, B. et al. Identification of ERGIC-53 as an intracellular transport receptor of α1-antitrypsin. J. Cell Biol. 180, 705–712 (2008).

9. Zhang, B. et al. Mice deficient in LMAN1 exhibit FV and FVIII deficiencies and liver accumulation of α1-antitrypsin. Blood 118, 3384–3391 (2011).

10. Zhu, M. et al. Analysis of MCFD2- and LMAN1-deficient mice demonstrates distinct functions in vivo. Blood Adv. 2, 1014–1021 (2018).

11. Vollenweider, F., Kappeler, F., Itin, C. & Hauri, H.-P. Mistargeting of the Lectin ERGIC-53 to the Endoplasmic Reticulum of HeLa Cells Impairs the Secretion of a Lysosomal Enzyme. J. Cell Biol. 142, 377–389 (1998).

12. Appenzeller, C., Andersson, H., Kappeler, F. & Hauri, H.-P. The lectin ERGIC-53 is a cargo transport receptor for glycoproteins. Nat. Cell Biol. 1, 330–334 (1999).

13. Chen, Y., Hojo, S., Matsumoto, N. & Yamamoto, K. Regulation of Mac-2BP secretion is mediated by its N-glycan binding to ERGIC-53. Glycobiology 23, 904–916 (2013).

14. Duellman, T., Burnett, J., Shin, A. & Yang, J. LMAN1 (ERGIC-53) is a potential carrier protein for matrix metalloproteinase-9 glycoprotein secretion. Biochem. Biophys. Res. Commun. 464, 685–691 (2015).

15. Anelli, T. et al. Sequential steps and checkpoints in the early exocytic compartment during secretory IgM biogenesis. EMBO J. 26, 4177–4188 (2007).

16. Zhang, B. Combined deficiency of factor V and factor VIII is due to mutations in either LMAN1 or MCFD2. Blood 107, 1903–1907 (2006).

17. Zhang, B. et al. Bleeding due to disruption of a cargo-specific ER-to-Golgi transport complex. Nat. Genet. 34, 6 (2003).

18. Kawasaki, N. et al. The sugar-binding ability of ERGIC-53 is enhanced by its interaction with MCFD2. Blood 111, 1972–1979 (2008).

19. Zheng, C. et al. Structural Characterization of Carbohydrate Binding by LMAN1 Protein Provides New Insight into the Endoplasmic Reticulum Export of Factors V (FV) and VIII (FVIII). J. Biol. Chem. 288, 20499–20509 (2013).

20. Guy, J. E., Wigren, E., Svärd, M., Härd, T. & Lindqvist, Y. New Insights into Multiple Coagulation Factor Deficiency from the Solution Structure of Human MCFD2. J. Mol. Biol. 381, 941–955 (2008).

21. Tempio, T. et al. A virtuous cycle operated by ERp44 and ERGIC-53 guarantees proteostasis in the early secretory compartment. iScience 24, 102244 (2021).

22. Fu, Y.-L., Zhang, B. & Mu, T.-W. LMAN1 (ERGIC-53) promotes trafficking of neuroreceptors. Biochem. Biophys. Res. Commun. 511, 356–362 (2019).

23. Huang, Y., et al. An in vitro vesicle formation assay reveals cargo clients and factors that mediate vesicular trafficking. Proc. Natl. Acad. Sci. 118, e2101287118 (2021).

24. Klaus, J. P. et al. The Intracellular Cargo Receptor ERGIC-53 Is Required for the Production of Infectious Arenavirus, Coronavirus, and Filovirus Particles. Cell Host Microbe 14, 522–534 (2013).

25. Zeyen, L., Döring, T. & Prange, R. Hepatitis B Virus Exploits ERGIC-53 in Conjunction with COPII to Exit Cells. Cells 9, 1889 (2020).

26. Kamiya, Y. Molecular Basis of Sugar Recognition by the Human L-type Lectins ERGIC-53, VIPL, and VIP36*. 283, 5 (2008).

27. Itin, C., Roche, A. C., Monsigny, M. & Hauri, H. P. ERGIC-53 is a functional mannose-selective and calcium-dependent human homologue of leguminous lectins. Mol. Biol. Cell 7, 483–493 (1996).

28. Itin, C., Schindler, R. & Hauri, H. P. Targeting of protein ERGIC-53 to the ER/ERGIC/cis-Golgi recycling pathway. J. Cell Biol. 131, 57–67 (1995).

29. Kappeler, F., Klopfenstein, D. R. C., Foguet, M., Paccaud, J.-P. & Hauri, H.-P. The Recycling of ERGIC-53 in the Early Secretory Pathway ERGIC-53 CARRIES A CYTOSOLIC ENDOPLASMIC RETICULUM-EXIT DETERMINANT INTERACTING WITH COPII. J. Biol. Chem. 272, 31801–31808 (1997).

30. Nufer, O. et al. Role of cytoplasmic C-terminal amino acids of membrane proteins in ER export. J. Cell Sci. 115, 619–628 (2002).

31. Lahtinen, U., Svensson, K. & Pettersson, R. F. Mapping of structural determinants for the oligomerization of p58, a lectin-like protein of the intermediate compartment and cis-Golgi. Eur. J. Biochem. 260, 392–397 (1999).

32. Schweizer, A., Fransen, J. A., Bächi, T., Ginsel, L. & Hauri, H. P. Identification, by a monoclonal antibody, of a 53-kD protein associated with a tubulo-vesicular compartment at the cis-side of the Golgi apparatus. J. Cell Biol. 107, 1643–1653 (1988).

33. Neve, E. P. A., Lahtinen, U. & Pettersson, R. F. Oligomerization and interacellular localization of the glycoprotein receptor ERGIC-53 is independent of disulfide bonds. J. Mol. Biol. 354, 556–568 (2005).

34. Satoh, T., Suzuki, K., Yamaguchi, T. & Kato, K. Structural Basis for Disparate Sugar-Binding Specificities in the Homologous Cargo Receptors ERGIC-53 and VIP36. PLoS ONE 9, e87963 (2014).

35. Satoh, T. et al. Crystallographic snapshots of the EF-hand protein MCFD2 complexed with the intracellular lectin ERGIC-53 involved in glycoprotein transport. Acta Crystallogr. Sect. F Struct. Biol. Commun. 76, 216–221 (2020).

36. Nishio, M. et al. Structural basis for the cooperative interplay between the two causative gene products of combined factor V and factor VIII deficiency. Proc. Natl. Acad. Sci. 107, 4034–4039 (2010).

37. Wigren, E., Bourhis, J.-M., Kursula, I., Guy, J. E. & Lindqvist, Y. Crystal structure of the LMAN1-CRD/MCFD2 transport receptor complex provides insight into combined deficiency of factor V and factor VIII. FEBS Lett. 584, 878–882 (2010).

38. Velloso, L. M., Svensson, K., Schneider, G., Pettersson, R. F. & Lindqvist, Y. Crystal Structure of the Carbohydrate Recognition Domain of p58/ERGIC-53, a Protein Involved in Glycoprotein Export from the Endoplasmic Reticulum. J. Biol. Chem. 277, 15979–15984 (2002).

39. Receveur-Bréchot, V. & Durand, D. How Random are Intrinsically Disordered Proteins? A Small Angle Scattering Perspective. Curr. Protein Pept. Sci. 13, 55–75 (2012).

40. Bepler, T. et al. Positive-unlabeled convolutional neural networks for particle picking in cryo-electron micrographs. Nat. Methods 16, 1153–1160 (2019).

41. Jumper, J. et al. Highly accurate protein structure prediction with AlphaFold. Nature 596, 583–589 (2021).

42. Punjani, A. & Fleet, D. J. 3D variability analysis: Resolving continuous flexibility and discrete heterogeneity from single particle cryo-EM. J. Struct. Biol. 213, 107702 (2021).

43. Yagi, H. et al. Improved secretion of glycoproteins using an N-glycan-restricted passport sequence tag recognized by cargo receptor. Nat. Commun. 11, (2020).

44. Lo, M. N., Damon, L. J., Wei Tay, J., Jia, S. & Palmer, A. E. Single cell analysis reveals multiple requirements for zinc in the mammalian cell cycle. eLife 9, (2020).

45. Yamada, T. et al. A novel missense mutation causing abnormal LMAN1 in a Japanese patient with combined deficiency of factor V and factor VIII. Am. J. Hematol. 84, 738–742 (2009).

46. Zhang, Y., Liu, Z. & Zhang, B. Separate roles of LMAN1 and MCFD2 in ER-to-Golgi trafficking of FV and FVIII. Blood Adv. 7, 1286–1296 (2023).

47. Zhang, Y. et al. LMAN1–MCFD2 complex is a cargo receptor for the ER-Golgi transport of α1-antitrypsin. Biochem. J. 479, 839–855 (2022).

48. Yerushalmi, N. et al. ERGL, a novel gene related to ERGIC-53 that is highly expressed in normal and neoplastic prostate and several other tissues. Gene 265, 55–60 (2001).

49. Nufer, O., Mitrovic, S. & Hauri, H.-P. Profile-based data base scanning for animal L-type lectins and characterization of VIPL, a novel VIP36-like endoplasmic reticulum protein. J. Biol. Chem. 278, 15886–15896 (2003).

50. Kowada, T. et al. Quantitative Imaging of Labile Zn2+ in the Golgi Apparatus Using a Localizable Small-Molecule Fluorescent Probe. Cell Chem. Biol. 27, 1521–1531.e8 (2020).

51. Amagai, Y. et al. Zinc homeostasis governed by Golgi-resident ZnT family members regulates ERp44-mediated proteostasis at the ER-Golgi interface. Nat. Commun. 14, 2683 (2023).

52. Liu, R. et al. Organelle-Level Labile Zn2+ Mapping Based on Targetable Fluorescent Sensors. ACS Sens. 7, 748–757 (2022).

53. Watanabe, S. et al. Zinc regulates ERp44-dependent protein quality control in the early secretory pathway. Nat. Commun. 10, 603 (2019).

54. Brown, G. D., Willment, J. A. & Whitehead, L. C-type lectins in immunity and homeostasis. Nat. Rev. Immunol. 18, 374–389 (2018).

55. Mnich, M. E., van Dalen, R. & van Sorge, N. M. C-Type Lectin Receptors in Host Defense Against Bacterial Pathogens. Front. Cell. Infect. Microbiol. 10, (2020).

56. Miller, M. H. et al. LMAN1 is a receptor for house dust mite allergens. Cell Rep. 42, 112208 (2023).

## Method reference

57. Takagi, H. et al. New high-brilliance small angle x-ray scattering beamline, BL-15A2 at the photon factory. AIP Conf. Proc. 2054, 060038 (2019).

58. Shimizu, N. et al. Software development for analysis of small-angle x-ray scattering data. AIP Conf. Proc. 1741, 050017 (2016).

59. Yonezawa, K., Takahashi, M., Yatabe, K., Nagatani, Y. & Shimizu, N. MOLASS: Software for automatic processing of matrix data obtained from small-angle X-ray scattering and UV-visible spectroscopy combined with size-exclusion chromatography. Biophys. Physicobiology 20, e200001 (2023).

60. Manalastas-Cantos, K. et al. ATSAS 3.0: expanded functionality and new tools for small-angle scattering data analysis. J. Appl. Crystallogr. 54, 343–355 (2021).

61. McCall, K. A. & Fierke, C. A. Colorimetric and fluorimetric assays to quantitate micromolar concentrations of transition metals. Anal. Biochem. 284, 307–315 (2000).

62. Mastronarde, D. N. Automated electron microscope tomography using robust prediction of specimen movements. J. Struct. Biol. 152, 36–51 (2005).

63. Snijder, J. et al. Vitrification after multiple rounds of sample application and blotting improves particle density on cryo-electron microscopy grids. J. Struct. Biol. 198, 38–42 (2017).

64. Kimanius, D., Dong, L., Sharov, G., Nakane, T. & Scheres, S. H. W. New tools for automated cryo-EM single-particle analysis in RELION-4.0. Biochem. J. 478, 4169–4185 (2021).

65. Zivanov, J. et al. New tools for automated high-resolution cryo-EM structure determination in RELION-3. eLife 7, e42166 (2018).

66. Punjani, A., Rubinstein, J. L., Fleet, D. J. & Brubaker, M. A. cryoSPARC: algorithms for rapid unsupervised cryo-EM structure determination. Nat. Methods 14, 290–296 (2017).

67. Asarnow, D., Palovcak, E. & Cheng, Y. asarnow/pyem: UCSF pyem v0.5. (2019) doi:10.5281/zenodo.3576630.

68. Punjani, A., Zhang, H. & Fleet, D. J. Non-uniform refinement: adaptive regularization improves single-particle cryo-EM reconstruction. Nat. Methods 17, 1214–1221 (2020).

69. Emsley, P., Lohkamp, B., Scott, W. G. & Cowtan, K. Features and development of Coot. Acta Crystallogr. D Biol. Crystallogr. 66, 486–501 (2010).

70. Yamashita, K., Palmer, C. M., Burnley, T. & Murshudov, G. N. Cryo-EM single-particle structure refinement and map calculation using Servalcat. Acta Crystallogr. Sect. Struct. Biol. 77, 1282–1291 (2021).

71. Afonine, P. V. et al. Real-space refinement in PHENIX for cryo-EM and crystallography. Acta Crystallogr. Sect. Struct. Biol. 74, 531–544 (2018).

72. Mirdita, M. et al. ColabFold: making protein folding accessible to all. Nat. Methods 19, 679–682 (2022).

73. Pettersen, E. F. et al. UCSF ChimeraX: Structure visualization for researchers, educators, and developers. Protein Sci. Publ. Protein Soc. 30, 70–82 (2021).

74. Waterhouse, A. M., Procter, J. B., Martin, D. M. A., Clamp, M. & Barton, G. J. Jalview Version 2--a multiple sequence alignment editor and analysis workbench. Bioinforma. Oxf. Engl. 25, 1189–1191 (2009).

